# Obesity-Driven Lung Lipidome Remodeling Suppresses NK Cell Activation and Antiviral Immunity to Influenza Infection

**DOI:** 10.64898/2026.03.06.710186

**Authors:** Pamela H. Brigleb, Matthew Frank, Lauren Rowland, Theresa Bub, Cliff Guy, Brandi Livingston, Alexandra Mandarano, Emily R. Sekera, John Bowling, Stacey Schultz-Cherry

## Abstract

Obesity is a major risk factor for severe influenza A virus (IAV) infection, however, the innate immune mechanisms underlying this increased vulnerability remain unclear. Here, we identify significant defects in natural killer (NK) cell antiviral responses in mice with diet-induced obesity. In lean mice, NK cells are critical for protection as NK cell depletion during IAV infection led to increased lung viral load, morbidity, and mortality. In contrast, in obese mice NK cell depletion had minimal impact on viral replication or survival. Notably, IAV infection in obese mice recapitulated the phenotype observed in NK cell-depleted lean mice, indicating that obesity is associated with preexisting NK cell dysfunction. Following IAV infection, obese NK cells in the lung were functionally impaired with diminished activation (CD69^+^), cytokine production (IFN-γ), and cytolytic activity (Granzyme B) accompanied by defects in the mTOR signaling pathway and reduced glycolytic and oxidative metabolism. Bulk and spatial lipidomics revealed obesity and infection-driven remodeling of the lung lipidome. We observed increased triglyceride accumulation, abundance of long-chain free fatty acids, and a shift toward monounsaturated phospholipid species, reshaping the lung microenvironment that coincides with NK cell metabolic dysfunction. Consistent with this lipid-rich environment, obese NK cells sustained high expression of the lipid transporter CD36 post-IAV infection and accumulation of intracellular lipids (LipidTOX^+^), consistent with mechanisms known to suppress NK cell function. Notably, short-term weight loss (4 weeks) was sufficient to restore NK cell metabolism, antiviral function, and survival following IAV infection. These findings uncover a lipid-associated mechanism regulating NK cell function and show it plays a critical role in defense against infection and that it is dysfunctional in obesity. We suggest that targeting immunometabolism could lead to new antiviral therapies and potentially improve vaccine efficacy, especially in high-risk populations such as obesity.

**GRAPHICAL ABSTRACT:** 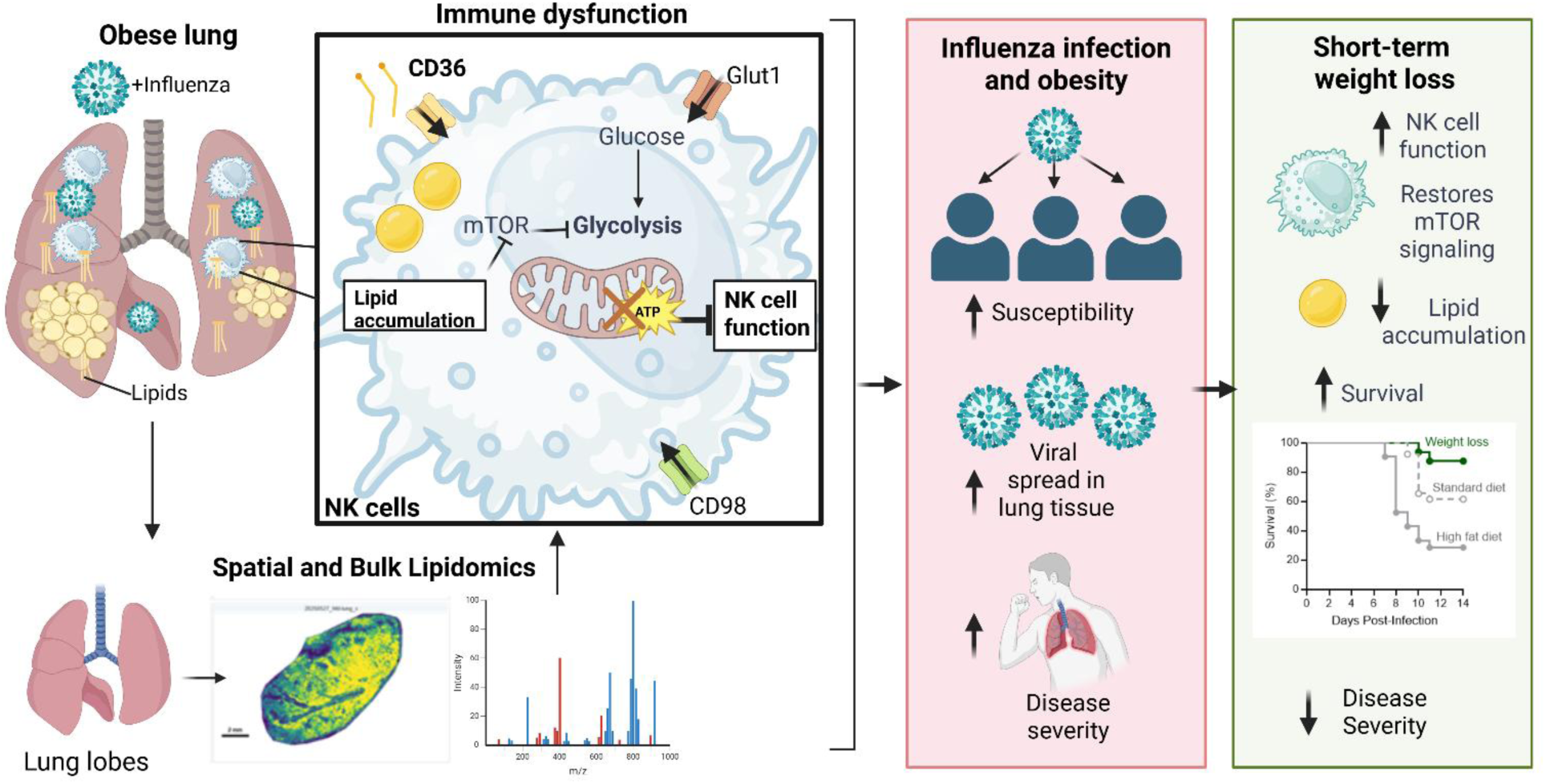

## INTRODUCTION

Influenza A virus (IAV) is an RNA virus that causes annual epidemics worldwide, resulting in approximately 1 billion infections and over half a million deaths each year^1,2^. Obesity and its associated metabolic disorders are a significant global health issue and are independent risk factors for severe IAV disease^3,4^. Clinical studies during the 2009 H1N1 pandemic identified obesity as a risk factor for hospitalization, mechanical ventilation, and death^5–7^. Obesity is associated with impaired antiviral immunity including blunted and delayed interferon (IFN) responses by the respiratory epithelium, resulting in greater early dissemination of virus throughout the lung^3,4,7–9^. However, the specific innate immune responses that contribute to insufficient early viral control and increased risk for IAV susceptibility and disease severity remain unclear.

IAV infects respiratory epithelial cells in the upper and lower respiratory tract, inducing cytokine production that recruits innate immune effector cells, including natural killer (NK) cells^1,10^. NK cells are antiviral lymphocytes that develop predominantly in the bone marrow^11–13^ and are rapidly recruited to the lung during IAV infection^10,14^. NK cells produce inflammatory cytokines, most prominently IFN-γ, and kill infected cells through cytolytic mechanisms^10,15^. NK cells contribute to antiviral responses in sublethal IAV infections in murine models and are important in the activation of adaptive immunity^14–17^. Individuals with genetic conditions that cause NK cell deficiencies such as GATA2 mutations have been reported to succumb to IAV infections^18,19^, highlighting their important antiviral role. Obesity disrupts systemic metabolic homeostasis and is associated with a reduction in circulating NK cells and diminished cytotoxic capacity, particularly in the context of cancer^17,18^. Emerging evidence suggests that immunometabolism is tightly linked to antiviral function^22–25^. However, it is not known whether and how obesity alters NK cell metabolic programming in the lung to compromise early antiviral defense, and whether these defects are reversible.

Herein, using a mouse model of diet-induced obesity (DIO), we demonstrate that obesity significantly impairs NK cell antiviral responses to IAV infection. Lung NK cells from obese mice have blunted activation, reduced effector function, and diminished metabolic flexibility including impaired mTOR-glycolytic signaling. Using bulk and spatial lipidomics, we reveal that obesity remodels the lung lipid landscape, including a shift toward monounsaturated phospholipid species that can affect cell membranes and mitochondrial activity. Obesity also causes triglyceride accumulation in the lung, and elevated lipid transporter expression in response to this lipid environment drives intracellular lipid accumulation in NK cells, which is known to impair NK cell function and metabolism. Notably, short-term weight loss reverses NK cell lipid accumulation, restores mTOR-dependent metabolic signaling, and rescues NK antiviral responses, ultimately improving mortality and morbidity following infection. Collectively, these findings reveal a lipid-associated mechanism of NK cell dysfunction in obesity during viral infection and highlight immunometabolism as a potential target to enhance antiviral immunity in vulnerable populations.

## RESULTS

### Obesity abrogates the protective role of NK cells during influenza infection

Obesity causes defects in early viral clearance resulting in increased viral spread in lungs. NK cells are important for the detection and clearance of IAV infected cells and production of inflammatory cytokines to amplify antiviral responses^10,15^. However, whether NK cells are protective or pathogenic in murine models depends on viral strain, dose, mouse background, and route of inoculation^10,15,26,27^. Furthermore, although obesity is associated with altered systemic NK immune responses, its impact on NK cell responses in the lung during IAV infection remains unclear. Since NK cells act within the first few days of infection, when obesity-associated defects in viral control and dissemination emerge, and because obesity alters NK cell biology in other contexts, we hypothesized that NK cells contribute to early viral control in lean mice but that this protection is compromised in obesity. To test this, we employed a DIO mouse model where WT C57BL/6 male mice were placed on standard diet (SD) or a 60% high fat diet (HFD) for a minimum of 12 weeks at which point HFD mice gained significantly more weight than SD mice and displayed signs of metabolic dysfunction^28^.

To test the contribution of NK cells to IAV infection in our model, SD or HFD mice were treated with either IgG2a isotype control or anti-NK1.1 (PK136) depleting antibody prior to and throughout infection (**Figure 1A, Figure S1A-C**). Mice were infected intranasally under light isoflurane anesthesia with a mild dose (10^3^ TCID_50_) of CA/09 H1N1 virus which resulted in 80% survival in our SD control mice (**Figure 1B**). NK depletion in SD mice led to significantly decreased survival of 25% (**Figure 1B**) and increased clinical scores (**Figure 1C**), with a trend toward greater weight loss (**Figure S1D**). This corresponded with higher lung viral titers at 3 dpi (**Figure 1D**) and increased pulmonary hemorrhaging (**Figure 1E**), indicating that NK cells are protective against IAV morbidity and mortality in SD lean mice in our model.

**Figure 1.**
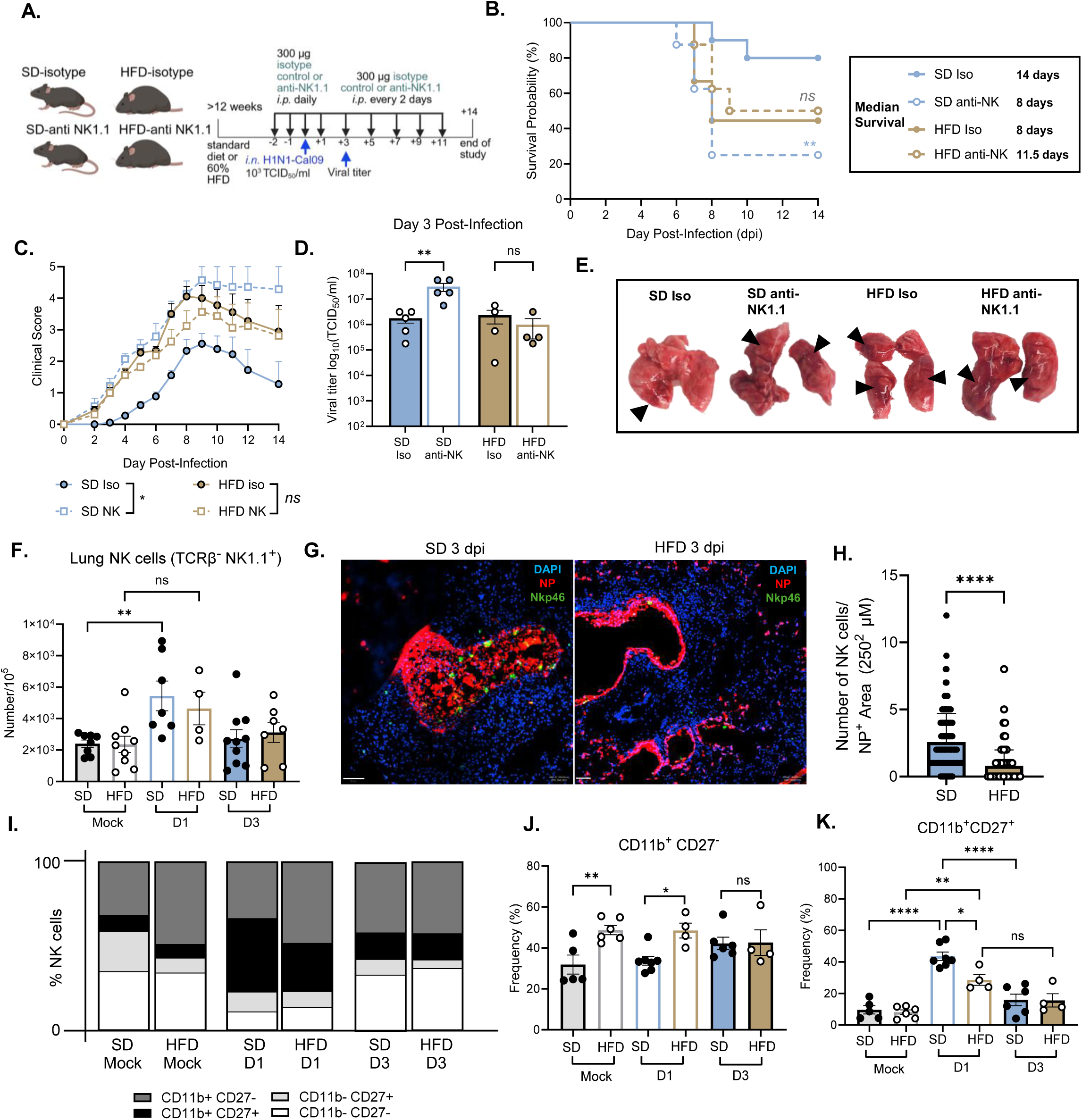
NK cell-mediated protection during IAV infection is abrogated in obesity. WT mice were placed on standard diet (SD) or 60% high-fat diet (HFD) for a minimum of 12 weeks. **(A-K)** WT and HFD mice were inoculated by intranasal (IN) infection with 1000 tissue culture infectious dose-50 (TCID_50_) of A/California/04/2009 (CA/09 H1N1) virus or PBS as a mock control. **(A-E)** NK cells were depleted with 300 μg of anti-NK1.1 PK136 antibody or an IgG2a isotype control days -2, -1, day of infection (d0), +1, and then every other day throughout the duration of the experiment. **(A)** Graphical summary and timeline of NK cell depletion studies made with *Biorender* **(B)** Probability of survival and median day survival time (n=9-10 per group) and **(C)** Clinical scores post-infection. **(D)** Lungs were harvested at 3 days-post infection (dpi) and viral titers were determined via TCID_50_ (*n=4-5* per group) **(E)** Representative gross lung images at 3 dpi with black arrows indicating areas of pulmonary hemorrhage. **(F, I-K)** At 1- or 3-dpi lungs were resected and processed for flow cytometry. Single-cell suspensions were stained for surface antibodies to characterize natural killer (NK) cells. **(F)** Number per 10^5^ lung cells of NK cells (CD45^+^, TCRβ^-^, NK1.1^+^) (*n=4-10* per group) **(G)** Lungs were processed for immunofluorescence at 3 dpi and stained for DAPI (blue), influenza NP protein (red) and NKp46 (NK cell marker, green) and **(H)** Number of NK cells per 250^2^ μm was quantified. **(I)** Average expression of the NK cell markers CD27 and CD11b to assess maturation stage. **(J)** Frequency of CD11b^+^ CD27^-^ lung NK cells and **(K)** Frequency of CD11b^+^CD27^+^ lung NK cells at baseline or at 1 or 3 dpi (*n=4-7* per group). Data are shown as mean ± SEM. Statistical significance was calculated using Mantel-Cox log-rank analysis **(A)** a two-way ANOVA **(C),** a One-way ANOVA with multiple comparison’s test **(D, F, J, K)** and a students *t* test with Welch’s correction **(H)**, * p < .05; ** p < .01; **** p < .0001; ns = not significant.

In contrast to SD, NK cell depletion in obese HFD mice had no significant impact on survival, viral replication, or clinical score (**Figure 1A-1E**). However, the obese HFD isotype group had a lower survival rate with increased clinical score and pulmonary hemorrhaging compared to the SD isotype group (**Figure 1B-E**). Notably, viral titers at 3 dpi were similar in SD and HFD isotype-treated groups, consistent with previous findings showing greater viral dissemination in obese lungs without significant differences in total lung titers^8^. Collectively, these data indicate NK cells are essential for protection during IAV infection in lean SD mice but fail to mediate protection in obese HFD mice, suggesting a functional impairment of NK cells in the obese lung.

### Obesity induces tissue-specific alterations in NK cell development and phenotype

To determine whether altered NK cell protection in obesity during IAV infection reflects defects in NK cell development or tissue-specific programming, we first examined NK cells at baseline. NK cells develop primarily in the bone marrow and circulate to distal tissues^11–13^. Although total immune cell (CD45^+^) and NK cell (CD45^+^ Tcrβ^-^ NK1.1^+^) numbers were comparable between SD and HFD mice in the bone marrow (**Figure S2A**), NK cells from obese mice exhibited significantly lower expression of transcription factors important for NK cell development, including T-bet and GATA3 and a trend toward reduced EOMES expression (**Figure S2A**). These data suggest that HFD-induced obesity disrupts NK cell developmental programming in the bone marrow.

Since obesity is characterized by chronic, low-grade inflammation, we next assessed NK cells in the spleen. While total immune and NK cell numbers were comparable between SD and HFD mice in the spleen, splenic NK cells from obese mice expressed higher levels of T-bet, consistent with heightened systemic inflammation in obesity (**Figure S2B**). Collectively, these data indicate that obesity differentially affects NK cell developmental and systemic programs at baseline.

We next evaluated lung NK cells during early IAV infection. Mice were infected with IAV and tissues were harvested at 1- and 3-days post-infection (dpi) to assess early innate immune responses. At baseline, NK cell numbers and frequencies (CD45^+^ Tcrβ^-^ NK1.1^+^) in the lung were comparable between SD and HFD mice (**Figure 1F and Figure S2C**). Both groups exhibited similar early increases in NK cell number (**Figure 1F**) and frequency (**Figure S2C**) in the lung following infection. However, quantitative image analysis of immunofluorescence staining of lung tissue revealed significantly fewer NK cells (NKp46^+^) within IAV nucleoprotein (NP^+^) lung regions in obese mice at 3 dpi, suggesting reduced NK cell representation within sites of active infection rather than a defect in overall recruitment to the lung (**Figure 1G & H**).

NK cell maturation shapes their effector potential and timing of responses during an infection. To determine whether maturation differences contribute to altered early responses in obesity, we assessed CD27 and CD11b expression on lung NK cells in SD and HFD mice infected with a mock control or IAV (**Figure S2D**). At baseline, HFD mice exhibited skewing toward terminally differentiated CD27^-^ CD11b^+^ NK cells and a reduction in early/intermediate CD27^+^ CD11b^-^ NK cells in the lung (**Figure 1I-K**). Early after infection (1 dpi), both groups transitioned toward more mature CD27^+^ CD11b^+^ NK cells; however, this transition was blunted in HFD mice, which had fewer CD27⁺CD11b⁺ cells compared to SD controls (**Figure 1I-K**). These findings indicate a delayed maturation of lung NK cells in obesity during acute infection, which may hinder timely antiviral responses.

Collectively, these data reveal that obesity alters NK cell development in the bone marrow, inflammatory programming in systemic tissues, and phenotypic and spatial responses in the lung during infection. These findings underscore the importance of considering tissue-specific NK cell alterations when evaluating impaired antiviral immunity in obesity.

### NK cell antiviral response to influenza infection is impaired in obesity

To determine how obesity alters NK cell responses during IAV infection, SD and HFD mice were inoculated intranasally with 10^3^ TCID_50_ CA/09 H1N1 or PBS as a mock control, and early NK cell responses were analyzed at 1 and 3 dpi. We first assessed transcription factor expression, as NK cell activation is characterized by downregulation of EOMES (EOMES^high^ to EOMES^intermediate^ ^(int)^) and upregulation of T-bet, which is essential for cytotoxicity and cytokine production^29,30^. Both SD and HFD mice had similar proportions of EOMES^int^ NK cells at baseline with a significant increase at 3 dpi (**Figure 2A & B**). However, induction of T-bet was delayed in HFD mice compared to SD controls (**Figure 2C & D**).

**Figure 2.**
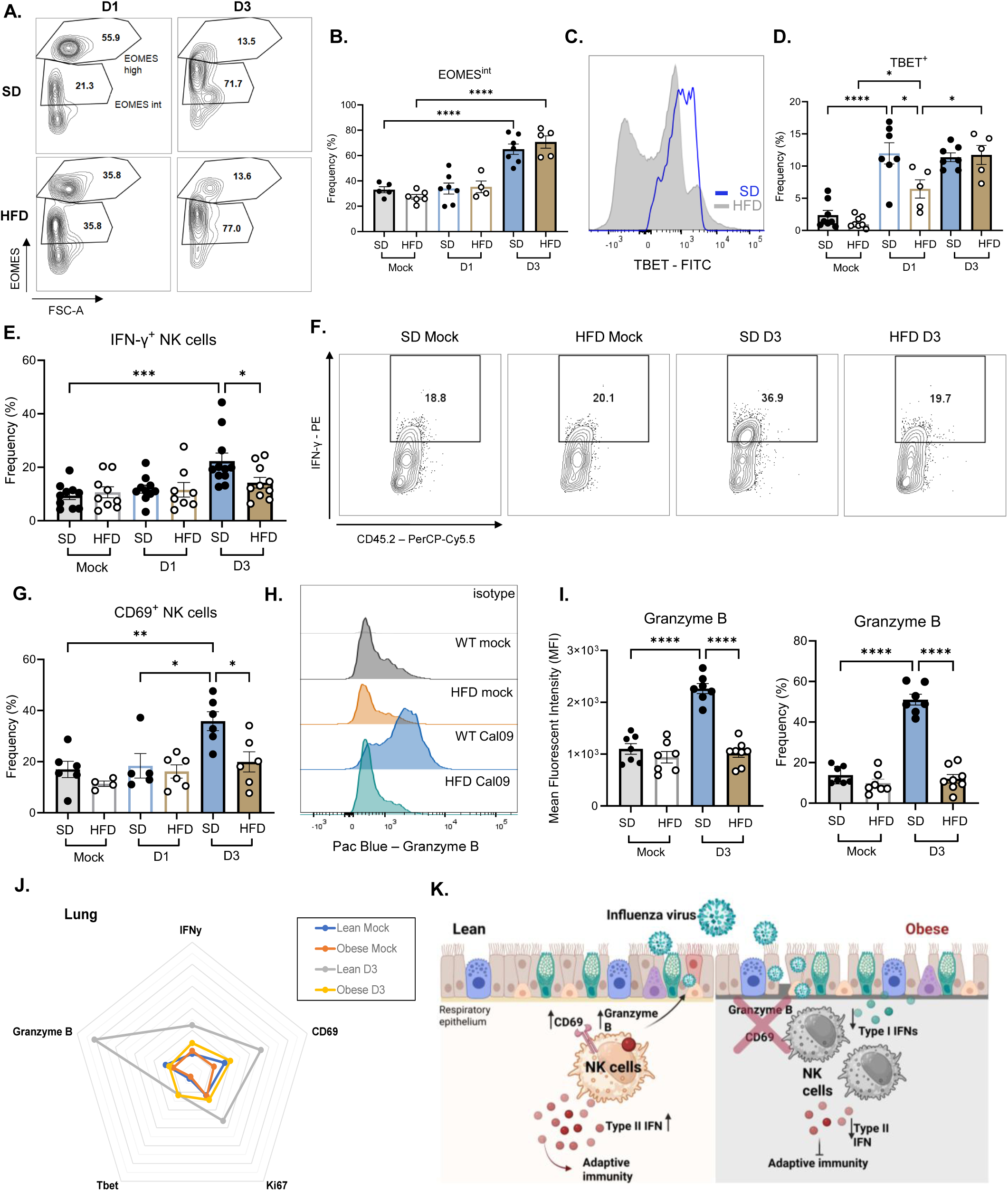
Obesity disrupts early transcriptional programming and sustained NK cell effector function post-IAV infection. WT mice were placed on SD or 60% HFD for a minimum of 12 weeks and were intranasally inoculated with 1000 TCID_50_ of CA/09 H1N1 virus or PBS (mock control) and lungs were resected and processed for flow cytometry at 1 or 3 dpi (n=4-11 per group). **(A)** Representative contour plots of EOMES expression in lung NK cells (CD45^+^, TCRβ^-^ NK1.1^+^) at 1 or 3 dpi **(B)** Frequency of intermediate EOMES expression in NK cells **(C)** Representative mean fluorescent intensity (MFI) plot normalized to mode of TBET expression in NK cells 1 dpi normalized to mode **(D)** Frequency of TBET expression in lung NK cells **(E)** Frequency of IFN-γ expression in NK cells **(F)** Representative contour plot of IFN-γ expression in NK cells in mock-infected controls or 3 dpi **(G)** Frequency of the early activation marker CD69 expression in NK cells **(H)** Representative MFI plot normalized to mode of Granzyme B expression and **(I)** MFI and frequency of Granzyme B expression in NK cells **(J)** IFNγ, Granzyme B, CD69, Tbet, and Ki67 expression displayed as spider plots. Data of NK cells from mock-infected WT mice in blue, mock-infected HFD mice in orange, WT 3 dpi in grey, and HFD 3 dpi in yellow. **(K)** Graphical summary of NK cell dysfunction in obesity during influenza infection, made with *Biorender*. Data are shown as mean ± SEM. Statistical significance was calculated using a one-way ANOVA with Tukey’s multiple comparison’s test (**B, D, E, G, & I)**. * p < .05; ** p < .01; *** p < .001; **** p < .0001; ns = not significant.

To assess whether these transcriptional and maturation differences translated into functional impairments, we assessed cytokine production (IFN-γ), early activation (CD69), and cytolytic activity (Granzyme B) in lung NK cells at baseline and early after infection. At baseline, there were no differences between SD and HFD groups (**Figure 2E-I**).

By 3 dpi, NK cells from SD mice mounted robust antiviral responses with significantly higher IFN-γ, CD69, and Granzyme B expression compared to baseline which are significantly blunted in NK cells from obese mice (**Figure 2E-I**). Although T-bet expression eventually reached levels comparable to SD mice by 3 dpi, its delayed induction coincided with impaired early functional responses. Overall, these results indicate that obesity delays NK cell transcriptional activation and disrupts the timely coordination of transcription factor induction with effector function, resulting in impaired NK cell antiviral responses during IAV infection (**Figure 2J & K**).

### Obesity uncouples nutrient transporter expression from mTOR activation in NK cells

Cellular metabolism is essential for immune cell function. NK cell metabolism influences their activation, proliferation, and function in response to viral infection^31^. Obesity is a metabolic disorder that alters nutrient availability and rewires metabolic pathways in immune cells^32–34^. Given the impaired NK cell effector responses observed in obesity, we next investigated whether defective metabolic programming contributes to dysfunctional NK cell antiviral immunity.

To assess NK cell metabolic activity during influenza infection, we first measured proliferation and cell size. NK cells typically proliferate and increase in size as they become more metabolically active^31^. At 3 dpi, lung NK cells from SD mice had increased Ki-67 expression (**Figure 3A**) and cell size (FSC-A) (**Figure 3B**), whereas NK cells from HFD mice did not, suggesting impaired metabolic activation in obesity.

**Figure 3.**
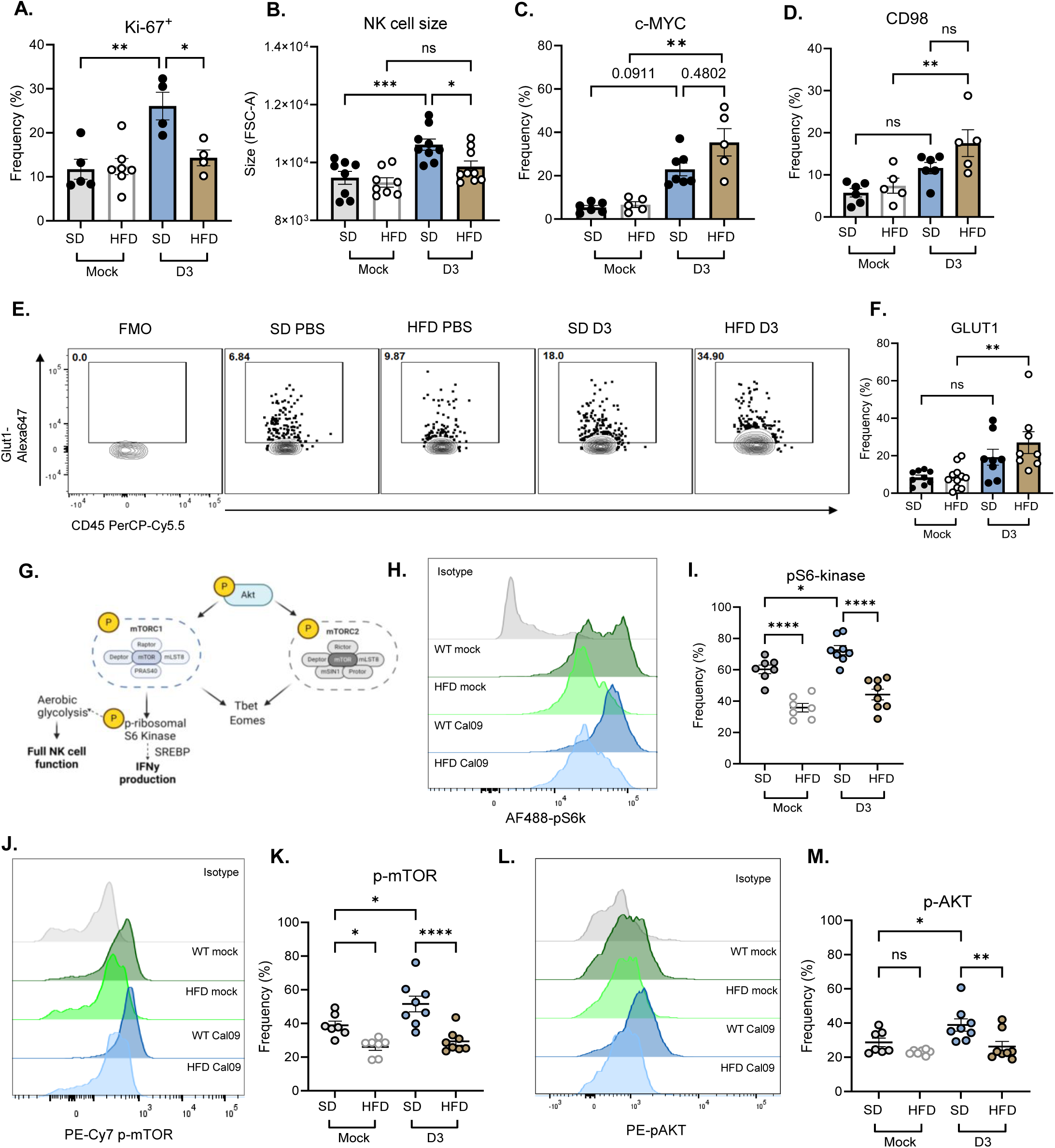
Metabolic uncoupling of nutrient transporter expression and mTOR activation in NK cells in obese mice. WT mice were placed on SD or 60% HFD for a minimum of 12 weeks and were intranasally inoculated with 1000 TCID_50_ of CA/09 H1N1 virus or PBS (mock control) and lungs were resected and processed for flow cytometry at 3 dpi to assess lung NK cells (CD45^+^, TCRβ^-^ NK1.1^+^). **(A)** Frequency of the marker of proliferation Ki-67 (n = 4-7/group) **(B)** Cell size as assessed by FSC-A (n = 8-9/group) **(C)** Frequency of c-MYC and **(D)** amino acid transporter CD98 expression (n=5-6/group) **(E)** Representative contour plots of GLUT1 expression on NK cells including fluorescence minus one control (FMO) and **(F)** Frequency of GLUT1 expression **(G)** Summary of mTOR signaling cascade and impact on NK cell function, made with *Biorender*. **(H)** Representative MFI plot normalized to mode of phosphorylated S6 kinase (pS6k) and **(I)** Frequency of pS6-kinase, **(J)** Representative MFI plot normalized to mode of phosphorylated mTOR and **(K)** Frequency of p-mTOR and **(L)** Representative MFI plot normalized to mode of phosphorylated AKT and **(M)** Frequency of p-AKT (n= 7-8/group). Data are shown as mean ± SEM. Statistical significance was calculated using a one-way ANOVA with Tukey’s multiple comparison’s test **(A-D, F, I, K, M)**. * p < .05; ** p < .01; *** p < .001; **** p < .0001; ns = not significant.

NK cells rely on several metabolites for maturation, differentiation, and effector functions, including glucose and amino acids^22–24,31,35–40^. We first examined these metabolic pathways by assessing nutrient transporter expression and c-Myc expression, which transcriptionally regulates the expression of nutrient transporters including the amino acid transporter CD98 and glucose transporter Glut-1^41–43^. c-Myc, CD98, and Glut-1 expression were modestly increased in SD NK cells following infection, whereas they were significantly upregulated in HFD NK cells (**Figure 3C-F**). This indicates that upstream induction of nutrient transporters and c-Myc pathways remain intact in obesity, with impairments instead occurring in downstream metabolic and functional processes.

Since early nutrient-sensing programs remained intact, we next examined downstream metabolic signaling pathways. Mammalian target of rapamycin (mTOR) and corresponding signaling pathways are a master regulator of metabolism that is required for NK cell function. Inhibition of mTOR with rapamycin impedes NK cell cytokine production through the activation of p-ribosomal S6 kinase (p-S6K) and other cytotoxic effector functions in both murine and human NK cells^44–46^. We assessed several key steps in the mTOR signaling pathway by phospho-flow cytometry, including p-AKT, p-mTOR, and p-S6K (**Figure 3G**). Of note, the p-mTOR antibody recognizes both mTORC1 and mTORC2 complexes. At baseline, there was reduced expression of p-mTOR and p-S6K in HFD NK cells but not p-AKT compared to SD controls, indicating that impairment of this signaling pathway may be predominately downstream of p-AKT (**Figure 3H-M**). However, these defects in mTOR signaling were more pronounced at 3 dpi, with increases in mTOR signaling in the SD compared to mock but not in the HFD NK cells (**Figure 3H-M**). These data identify a metabolic uncoupling in obesity, wherein NK cells induce nutrient transporters to acquire energy needed for effector functions but fail to activate downstream mTOR signaling which is required for antiviral responses.

### Glycolytic and mitochondrial defects in obese NK cells post-IAV infection

As mTOR integrates nutrient signals to support mitochondrial respiration and glycolysis, we next examined whether the defects we observed in the mTOR-pS6K pathway were accompanied by impaired metabolic function in obese NK cells post-IAV infection. Glycolysis and mitochondrial function are essential for NK cell effector function^40,47,48^. At 3 dpi, lung NK cells from SD or HFD mice were assessed for glycolysis measured by extracellular acidification rate (ECAR) and proton efflux rate (PER) and for mitochondrial respiration by measuring oxygen consumption rate (OCR) (**Figure 4A**). HFD NK cells had markedly reduced metabolic activity compared to SD NK cells post-IAV infection, as shown by reduced OCR, ECAR, and PER curves (**Figure 4B-D**). HFD NK cells had significantly reduced basal and ATP-linked respiration and failed to increase mitochondrial activity in response to metabolic stress, resulting in a lower maximal respiration and spare respiratory capacity (**Figure 4E-G**). There was less proton leakage in HFD NK cells, which is likely due to their overall low mitochondrial activity (**Figure 4H**) and the coupling efficiency remained comparable between SD and HFD NK cells (**Figure 4I**). Collectively, these results indicate that the mitochondrial impairment observed in HFD NK cells may be due to a failure to activate metabolic pathways rather than increased membrane leakage.

**Figure 4.**
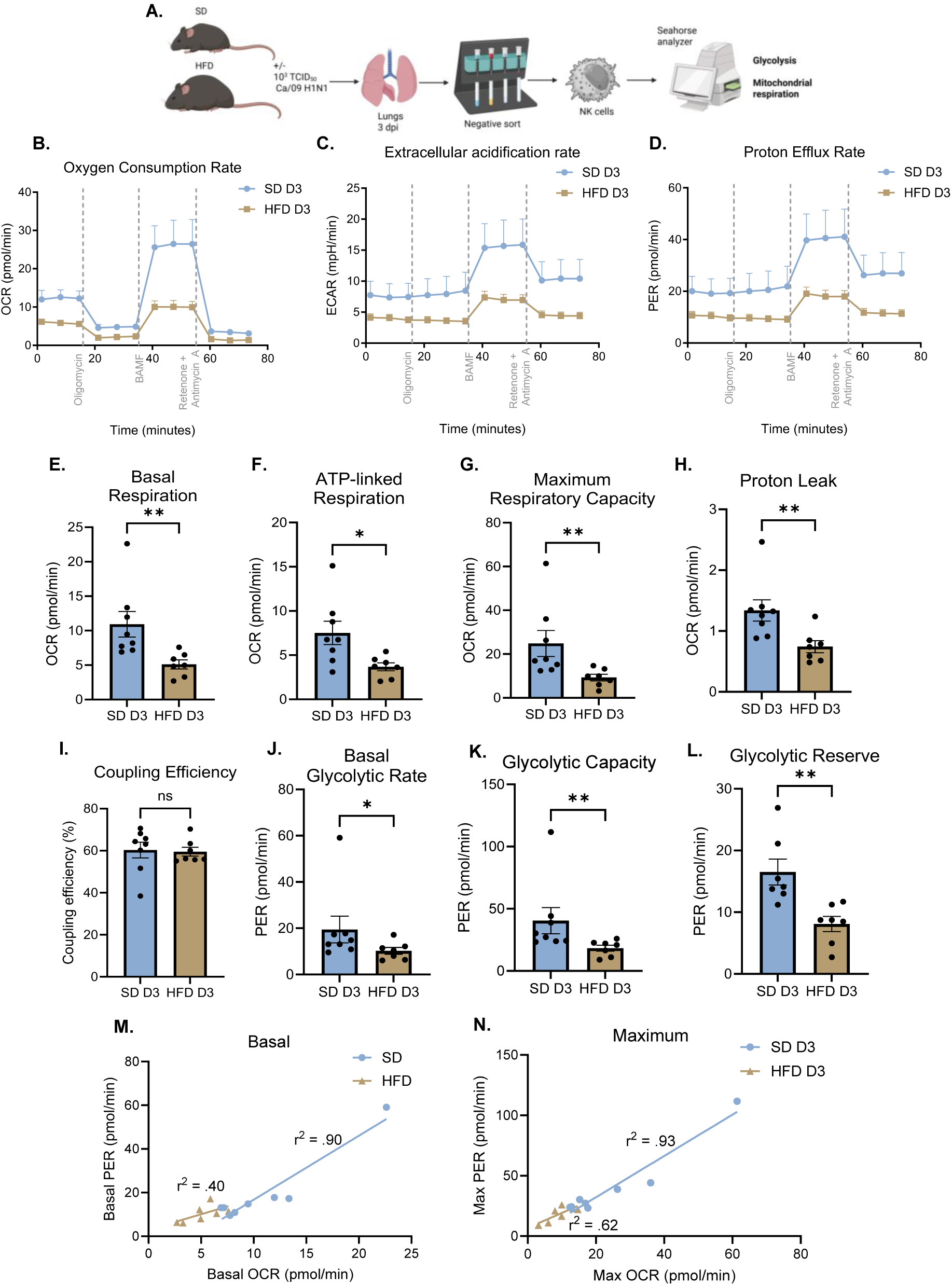
Obesity impairs lung NK cell metabolic flexibility during influenza infection. WT mice were placed on SD or 60% HFD for a minimum of 12 weeks and were intranasally inoculated with 1000 TCID_50_ of CA/09 H1N1 virus. At 3 dpi, lungs were resected and processed for single cell sorting to isolate lung NK cells by magnetic activated cell sorting for Seahorse metabolic analysis (n=7-8/group). **(A)** Summary of workflow made with *Biorender* **(B)** Oxygen consumption rate (OCR) **(C)** extracellular acidification rate (ECAR) and **(D)** proton efflux rate (PER) over time. **(E-I)** Metabolic analyses based on **B** including basal respiration **(E)**, ATP-linked respiration **(F)**, maximum respiratory capacity **(G)**, proton leak **(H)** and coupling efficiency **(I). (J-L)** Metabolic analyses based on **D** including basal glycolytic rate **(J)**, glycolytic capacity **(K)**, and glycolytic reserve **(L). (M, N)** Linear regression correlation analysis of basal PER vs. basal OCR **(M)** or maximum PER vs. maximum OCR **(N)**. Data are shown as mean ± SEM. Statistical analyses include a students *t* test with Welch’s correction **(E-L)**. * p < .05; ** p < .01; ns = not significant.

HFD NK cell glycolytic responses were similarly blunted, with reduced basal glycolytic rate, capacity and reserve (**Figure 4J-L**). These findings indicate that despite the upregulation of Glut-1 post-IAV infection in HFD NK cells, they fail to utilize glucose as energy for glycolysis. This could be explained by the defects in the mTOR-pS6K we observed (**Figure 3**). We next assessed whether NK cells coordinated mitochondrial and glycolytic activity during IAV infection. In SD mice, OCR and PER were positively correlated, indicated a flexible and coordinated metabolic response to a viral infection (**Figure 4M & N**). However, this correlation was disrupted in HFD NK cells, where there was not a strong correlation between mitochondrial respiration and glycolytic activity (**Figure 4M & N**). This uncoupling supports that obese NK cells are metabolically inflexible and are unable to execute these metabolic programs required for full antiviral effector function.

### Obesity and IAV infection remodel the lung lipidome

Diet-induced obesity in mice causes lipid accumulation and metabolic injury in the liver^28^, but how obesity alters the lung lipid environment during IAV infection is poorly understood. Lipids shape lung physiology by regulating membrane structure and fluidity, pulmonary surfactant, mitochondrial function, and immune signaling pathways relevant to NK cell activation^49,50^. Given the metabolic defects observed in obese NK cells, we investigated whether obesity and IAV infection remodel the lung lipidome in ways that could influence antiviral immunity.

To characterize obesity and infection-driven lipid remodeling, lungs from SD and HFD mice inoculated with PBS (mock) or CA/09 were collected at 3 dpi and analyzed using complementary bulk and spatial lipidomics workflows (**Figure 5A**). Bulk lipidomics of whole-lung homogenates provided representative lung-wide lipid profiles and assessed major phospholipid classes, including phosphatidylcholine (PC; abundant in cell membranes and pulmonary surfactant), phosphatidylglycerol (PG; enriched in mitochondrial and surfactant membranes), and phosphatidylethanolamine (PE; important for membrane curvature and mitochondrial integrity) (**Figure S3A-C**). Spatial lipidomics (matrix assisted laser desorption ionization mass spectrometry imaging, MALDI-MSI) was conducted on lung sections, with adjacent sections stained for DAPI and IAV NP to distinguish infected from uninfected regions. Together, these approaches enabled assessment of both global lipid composition and spatial remodeling in response to obesity and IAV infection.

**Figure 5.**
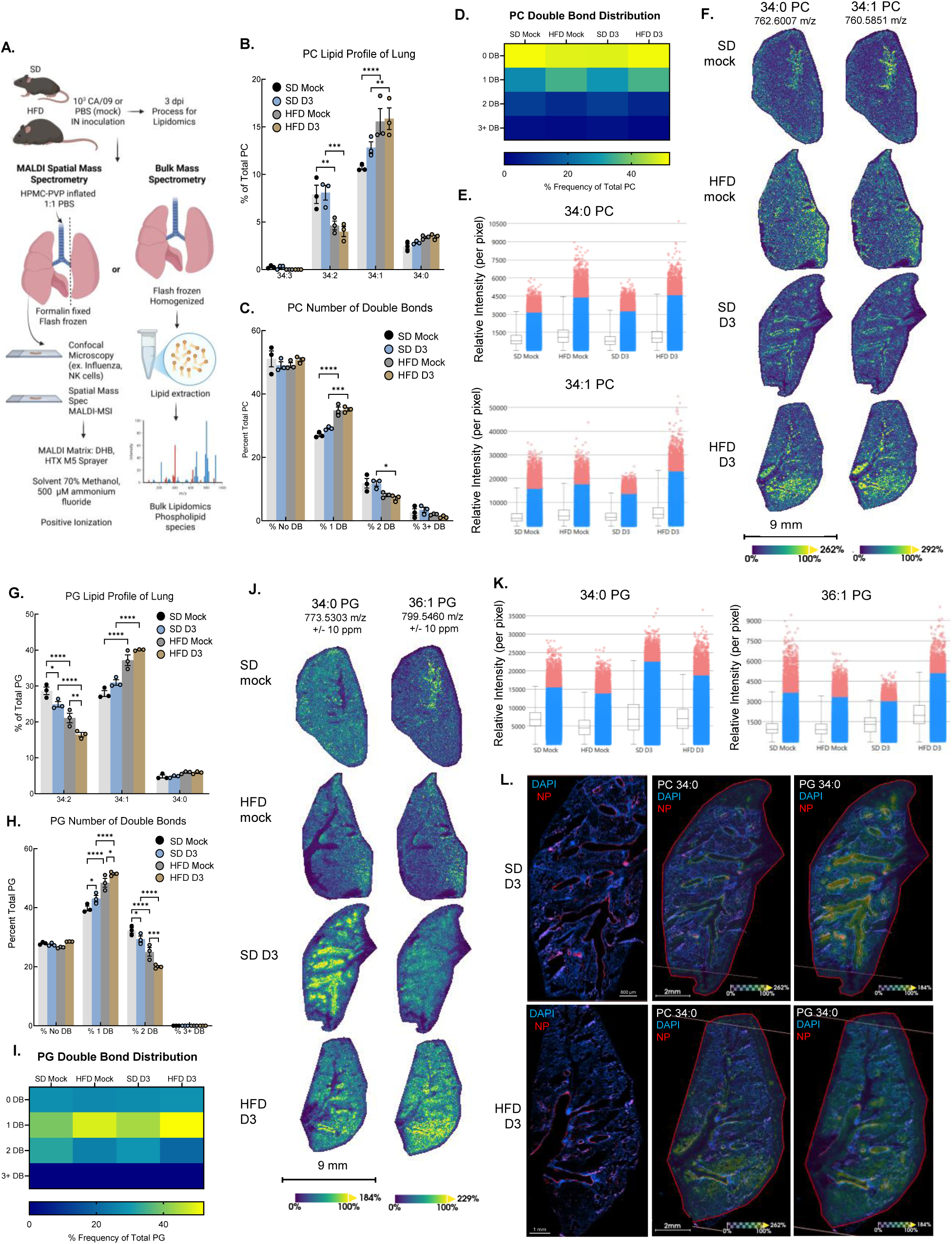
Obesity and influenza infection remodels the lung lipidome. WT mice were placed on SD or 60% HFD for a minimum of 12 weeks and were intranasally inoculated with 1000 TCID_50_ of CA/09 H1N1 virus or PBS (mock control). At 3 dpi, lungs were resected and processed for either bulk or spatial lipidomics. **(A)** Summary of workflow of sample processing made with *Biorender* **(B-D, G-I)** Bulk lipidomic analyses of relative abundance of phospholipid species in the lung. **(B)** Representative relative abundance of 34-carbon length PC lipids **(C)** Relative percentage of number of double bond distribution among all analyzed PC lipids and summarized in a heat map plot in **(D). (E & F)** MALDI spatial mass spectrometry including relative intensity per pixel of 34:0 PC (762.6007 +/- 10 ppm) or 34:1 PC (760.5851 +/- 10 ppm) **(E)** and distribution of 34:0 or 34:1 PC in lung tissue normalized by total ion current (TIC). **(G)** Representative relative abundance of 34-carbon length PG lipids **(H)** Relative percentage of number of double bond distribution among all analyzed PG lipids and summarized in a heat map plot in **(I). (J & K)** MALDI spatial mass spectrometry including distribution of 34:0 or 36:1 PG in lung tissue normalized by total ion current (TIC) **(J)** and relative intensity per pixel of 34:0 PG (773.5303 +/- 10 ppm) or 36:1 PG (799.5460 +/- 10 ppm). **(L)** Overlay of serial lung sections stained by IF with DAPI (blue) or influenza nucleoprotein (NP) indicating sites of active viral replication (red) with either 34:0 PC or PG. Data are shown as mean ± SEM. Statistical significance was calculated using a two-way ANOVA with multiple comparison’s **(B, C, G, H)**. * p < .05; ** p < .01; *** p < .001; **** p < .0001; ns = not significant.

We first examined PC, the most abundant lung phospholipid^50,51^. Since the degree of PC which contains unsaturated fatty acids influences membrane organization, mitochondrial function, and immune signaling, we analyzed individual PC species and the distribution of double bonds in the fatty acids across the PC lipid pool. We focused on the 34-carbon PC pool, which were among the most abundant species and were consistently detected in both bulk and spatial lipidomics datasets. Bulk lipidomics revealed minimal changes to PC 34:0 which contains 2 saturated fatty acids with obesity or infection; however, PC 34:1 which contains one fatty acid with one double bond and one saturated fatty acid increased with infection in SD lungs and were already elevated at baseline in HFD lungs, indicating that infection-induced remodeling in SD lungs phenocopies the baseline lipid composition of obese lungs (**Figure 5B**). In contrast, PC 34:2 and 34:3 were reduced in obese lungs, resulting in a shift toward fewer double bonds in the fatty acids across this PC lipid pool (**Figure 5B**), consistent with stress-associated lipid remodeling^52,53^. Similar patterns were observed across other abundant PC species, including 32- and 36-carbon length PCs (**Figure 5C & D, Figure S4A**), indicating broad obesity-driven remodeling of pulmonary PC composition. Spatial lipidomics further revealed marked differences in the distribution of PC species. Although the relative abundance of PC 34:0 did not significantly change as assessed by bulk lipidomics or spatial intensity analysis (**Figure 5B, E**), its regional localization was altered. Infected SD lungs displayed focal enrichment of PC 34:0 (and to a lesser extent, PC 34:1), whereas HFD lungs exhibited a more diffuse distribution with blunted regional remodeling (**Figure 5F**).

We next assessed PG, which is enriched in mitochondrial membranes and is the precursor for cardiolipin which is particularly relevant given the impaired mitochondrial respiration observed in obese NK cells^54^ (**Figure 4**). The 34-carbon pool of PG showed the same trends in obesity and IAV infection as the 34-carbon PC pool with increased PG 34:1 and decreased 34:2 (**Figure 5G**), and this trend was observed across the total PG lipid pool (**Figure 5H & I, Figure S4B**). Comparable shifts toward increased lipid pools containing a fatty acid with one double bond were observed in PE species (**Figure S4C & D**). As with PC, saturated PG 34:0 abundance was not significantly altered by obesity or infection, but its spatial distribution was re-modeled. Infection-associated regional enrichment of PG 34:0 was observed in SD but not HFD lungs (**Figure 5J & K**), and PG species containing a fatty acid with one double bond (ex. PG 36:1) exhibited increased spatial distribution and relative intensity in HFD lungs during IAV infection (**Figure 5J & K**).

Since spatial lipid remodeling appeared localized during infection, we examined adjacent lung sections stained for IAV NP to determine whether lipid redistribution correlated with sites of viral replication. NP staining confirmed that PC and PG 34:0 enrichment localized to regions of active viral infection (NP^+^) in SD but not HFD lungs. In SD mice, lipid enrichment was spatially restricted to airway- and parenchyma-associated infected areas, whereas obese lungs exhibited diffuse lipid distribution independent of viral localization (**Figure 5L**). These results indicate that obesity disrupts infection-driven lipid reorganization.

Collectively, these data demonstrate that obesity remodels the lung phospholipid landscape, characterized by increased monounsaturated and reduced polyunsaturated phospholipids, and impairs spatial lipid reorganization in response to viral infection. These changes are known to influence membrane properties, mitochondrial function, and metabolic signaling, providing a plausible mechanism by which obesity alters antiviral immunity, including NK cell responses.

### Obesity promotes a triglyceride-rich lung environment and enhances lipid uptake and accumulation in NK cells during IAV infection

Lipid metabolism is essential for immune cell function and homeostasis^25,55^, whereas excessive intracellular lipids can impair immune effector functions^56–59^. Prior studies demonstrated that exposure of NK cells to long-chain fatty acids (LCFAs), including palmitic and stearic acid, can suppress cytotoxicity, cytokine production, and cancer surveillance ^58,59^. Given the obesity-driven lung lipid remodeling and NK cell dysfunction observed, we hypothesized that the obese lung microenvironment promotes lipid exposure and accumulation in NK cells during IAV infection.

Triglycerides, which are precursors to LCFA generation, were significantly elevated in the HFD lung with ∼30-fold higher per mg of lung tissue compared to SD controls both at baseline and during IAV infection (**Figure 6A**). Spatial lipidomics revealed widespread triglyceride deposition throughout HFD lungs (**Figure 6B & C**), indicating that obesity establishes a triglyceride-enriched pulmonary environment in addition to changes in circulating lipids. Consistent with this finding, long chain fatty acids (LCFAs), including palmitoleic, palmitic, oleic, and stearic acid, were elevated in the obese lung at baseline compared to SD controls, indicating HFD-mediated lung lipid remodeling occurring prior to infection (**Figure 6D**). Following IAV infection, LCFAs increased in SD, indicating that IAV infection induces the release of free fatty acids (**Figure 6D**) to levels comparable to the levels that already existed in the HFD mice at baseline for palmitoleic, palmitic, and stearic acid. In the obese mice, LCFAs levels sustained high expression post-IAV infection. Interestingly, oleic acid had significantly higher abundance both at baseline and post-IAV infection in HFD lungs compared to SD controls. Oleic acid, in particular with the addition to palmitic acid, has been shown to inhibit NK cell function and mTOR signaling^59^.

**Figure 6.**
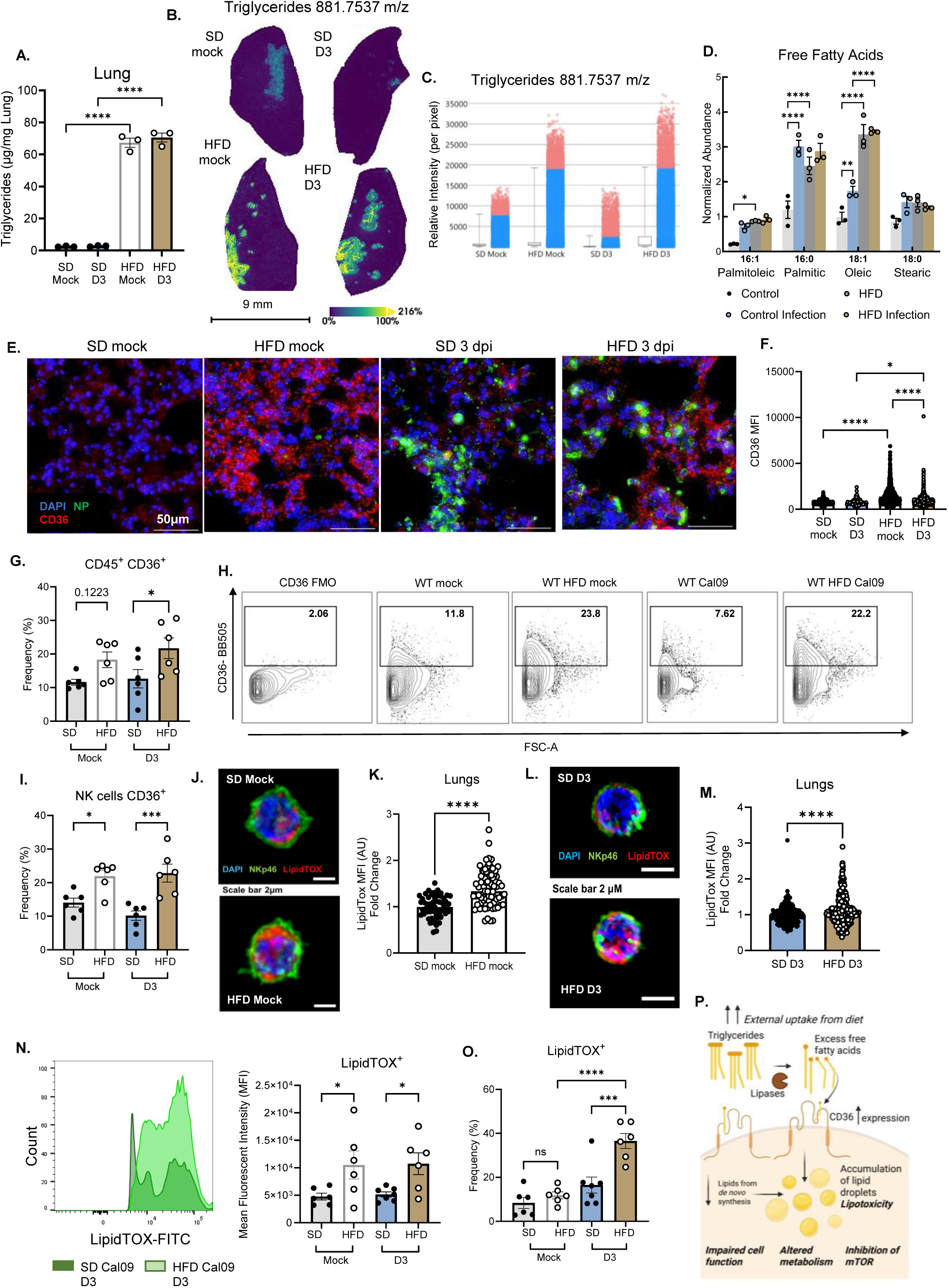
The obese lung microenvironment drives NK cell CD36 upregulation and lipid accumulation. WT mice were placed on SD or 60% HFD for a minimum of 12 weeks and were intranasally inoculated with 1000 TCID_50_ of CA/09 H1N1 virus or PBS (mock control). At 3 dpi, lungs were resected and processed for various analyses. **(A)** Triglyceride levels in lung tissue (n= 3/group) or **(B)** Triglyceride species (881.7537 m/z +/- 10 ppm) visualized in lung tissue by MALDI spatial mass spectrometry normalized by TIC and relative intensity per pixel was quantified in **(C)**. **(D)** Normalized abundance of long chain fatty acids in lung tissue. **(E & F)** Lung tissue was processed for IF at 3 dpi **(E)** Representative tissue sections stained for DAPI (blue), influenza NP protein for active viral replication (green) or the lipid transporter CD36 (red) and **(F)** CD36 MFI was determined with multiple fields per lung for at least 2 mouse lungs per group (>= 112 fields analyzed). **(G-I)** Lungs were resected and processed for flow cytometry at 3 dpi (n=6/group) to assess CD36 expression on **(G)** total immune cells (CD45^+^) **(H)** Representative contour plots of CD36 expression on lung NK cells (CD45^+^, TCRβ^-^ NK1.1^+^) including an FMO control and **(I)** frequency of CD36 expression. **(J-M)** At 3 dpi with IAV or PBS (mock), lungs were resected and processed for single cell sorting to isolate lung NK cells by magnetic activated cell sorting for IF imaging of NK cells (NKp46+, green), DAPI (blue) and LipidTOX (red) **(J & L)** Representative images with **(K & M)** MFI quantitation of LipidTOX staining during mock infection or 3 dpi with IAV (n>+ 178 cells/group analyzed). **(N, O)** LipidTOX was also assessed by flow cytometry (n= 6-7/group) with **(N)** representative MFI plot normalized to mode with MFI expression and **(O)** Frequency of NK cells expressing LipidTOX. **(P)** Graphical summary of lipid driven mechanisms of NK cell dysfunction made with *Biorender*. Data are shown as mean ± SEM. Statistical significance was calculated using a students *t* test with Welch’s correction **(A, K, M)**, or a One-way ANOVA with Tukey’s multiple comparison’s test **(B, F, G, I, N, O)**. * p < .05; *** p < .001; **** p < .0001; ns = not significant.

Triglyceride-derived LCFAs are taken up by lipid transporters including CD36. We assessed CD36 expression in lung tissue, and immunofluorescence revealed a significant increase in CD36 expression in HFD lungs at baseline and at 3 dpi compared to SD controls (**Figure 6E & F**). A reduction in CD36 in HFD IAV relative to HFD mock was observed, which may reflect infection-associated tissue injury and warrants further investigation.

NK cells express lipid transporters including CD36, and high expression of CD36 on NK cells has been linked to NK cell dysfunction and impaired tumor-killing^59^. We found that both total lung immune cells and lung NK cells have increased CD36 expression in HFD mice compared to SD mice, and that his expression doesn’t change following IAV infection at 3 dpi (**Figure 6G-I, Figure S5A**). This indicates there could be increased trafficking of LCFAs into lung NK cells in HFD mice through CD36 and other lipid transporters.

We next assessed whether increased lipid availability and transporter expression corresponded with intracellular lipid accumulation. We used LipidTOX which fluorescently labels intracellular neutral lipids and has a high affinity for lipid droplets. LipidTOX staining revealed significantly greater neutral lipid content in NK cells from HFD lungs at baseline and 3 dpi by both confocal microscopy and flow cytometry (**Figure 6J-O**). An increase in LipidTOX MFI and a higher frequency of LipidTOX^+^ NK cells was observed (**Figure 6N-O**). This increase was not detected in splenic NK cells, indicating that lipid accumulation in NK cells is tissue specific (**Figure S5E & F**). Similar trends were observed in total immune cells (CD45^+^) and NKT cells but not observed in conventional T cells (Tcrβ^+^) or NK^-^ Tcrβ^-^ immune populations (**Figure S5B**), consistent with preserved IFN-γ responses in these populations and only modest impairment in NKT cells (**Figure S5C**). Although CD36 expression also increased in other immune cell subsets post-infection (**Figure S5D**), the most pronounced and functionally relevant changes occurred in NK cells.

Collectively, these data demonstrate that obesity generates triglyceride and LCFA-rich lung environment and is associated with upregulation of lipid uptake programs including CD36, which is associated with increased intracellular lipid accumulation in NK cells. Given prior evidence that lipid accumulation suppress mTOR signaling and NK cell effector functions^57–59^, these findings position lipid exposure as a physiologically relevant contributor to NK cell dysfunction during IAV infection in obesity (**Figure 6P**).

### Short-term weight loss restores NK cell function and metabolic programming, improving protection against IAV infection in formerly obese mice

We previously found that short-term weight loss (4 weeks) in formerly obese mice was sufficient to restore metabolic function of splenic T cells and rescues vaccine recall responses^28^. We next asked whether obesity-associated NK cell dysfunction and metabolic rewiring were permanent or reversible, and if short-term weight loss could similarly improve outcomes following primary IAV infection. To test this, mice were placed on either SD or HFD for 16 weeks, after which a subset of HFD mice were diet-switched to SD for an additional 4 weeks prior to IAV infection (**Figure 7A**). Diet-switched mice lost weight during this period (**Figure 7B**) and had decreased total cholesterol, HDL, and LDL compared to the HFD controls (**Figure 7C-E**). Systemic metabolic improvements were also reflected in normalization of liver discoloration consistent with lipid accumulation (**Figure 7F**), reduced total liver weight (**Figure 7G**), and reduced pleural fat deposits surrounding the lung (**Figure 7H**). These data confirm that short-term diet-switch results in weight loss and improves systemic metabolic health of formerly obese mice.

**Figure 7.**
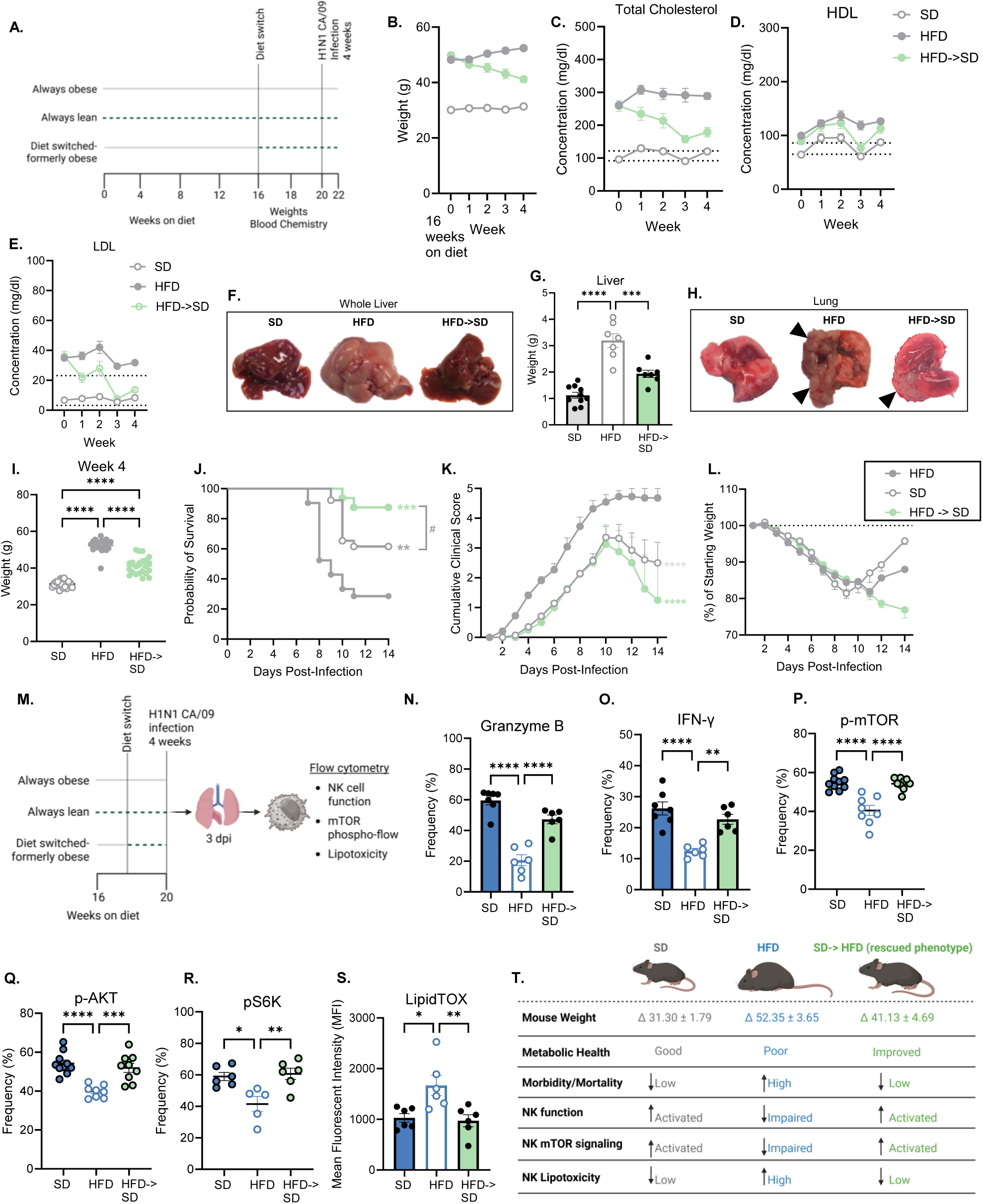
Weight loss restores NK cell function, mTOR signaling, and protection against influenza in formerly obese mice. **(A)** Timeline of diet and infection made with *Biorender* **(B)** Weights over time at start of diet-switch (HFD->SD) for all groups measured weekly **(C)** Total cholesterol **(D)** high-density lipoprotein (HDL) and **(E)** low-density lipoprotein (LDL) in blood measured weekly post-diet switch (n=11-15/group). **(F)** Representative images of gross anatomy livers, **(G)** liver weights (n=7-10/group) and **(H)** representative images of gross anatomy lungs with fatty deposits (black arrows) at 4 weeks post-diet switch. **(I)** Mouse weight at week 4 post-diet switch including SD and HFD controls at time of IAV infection. **(J-T)** Mice were intranasally inoculated with 1000 TCID_50_ of CA/09 H1N1 virus and assessed (n=16-26/group) for **(J)** survival **(K)** clinical scores and **(L)** weight loss, with the diet-switch group weight loss adjusted for weight loss only without infection average weekly weight loss **(M)** Summary of NK cell characterization by flow cytometry made in *Biorender* **(N-T)** At 3dpi, lungs were resected and processed for flow cytometry to assess lung NK cells (CD45^+^, TCRβ^-^ NK1.1^+^) (n= 6-7/group) including frequency of **(N)** Granzyme B **(O)** IFN-γ **(P)** phosphorylated mTOR **(Q)** phosphorylated AKT **(R)** phosphorylated S6 kinase and **(S)** MFI of intracellular LipidTOX staining. **(T)** Graphical summary of results from short term weight loss experiment and impact on NK cell function made with *Biorender*. Data are shown as mean ± SEM. Statistical significance was calculated a One-way ANOVA with Tukey’s multiple comparison’s test **(G, I, N-S)**, a Mantel-Cox log-rank analysis **(J)** and a two-way ANOVA **(K & L).** * p < .05; ** p < .01; *** p < .001; **** p < .0001.

We next evaluated whether short-term weight loss was sufficient to improve protection against primary IAV infection. At the time of infection, diet-switched mice were still significantly heavier than SD controls despite the improvements in metabolic health observed (**Figure 7I**). Consistent with our prior findings, HFD mice had severe morbidity and a low survival rate, whereas SD mice had a ∼60% survival rate and moderate disease (**Figure 7J-I**). Strikingly, weight-loss mice had significantly improved survival compared to the HFD controls, with a >80% survival rate and reduced morbidity (**Figure 7J-I**). Survival in the weight-loss group also trended higher than SD controls (p=.06), suggesting potential enhanced resistance warranting further studies (**Figure 7J**).

To determine whether the improved outcomes in the weight-loss group coincided with restoration of NK cell function and metabolism, lungs were harvested at 3 dpi and processed for flow cytometry to assess NK function, the mTOR pathway activation, and intracellular lipid accumulation (**Figure 7M**). Short-term weight loss significantly improved NK cell function including cytolytic activity (Granzyme B) and cytokine production (IFN-γ) compared to HFD controls (**Figure 7N, O**). Weight loss also restored mTOR signaling (p-mTOR, p-AKT, p-S6K) that were suppressed in obese NK cells (**Figure 7P-R**), and this recovery coincided with reduced intracellular neutral lipid accumulation as measured by LipidTOX (**Figure 7S**).

In summary, short-term weight loss reverses obesity-induced NK cell functional and metabolic defects and significantly improves protection against IAV infection (**Figure 7T**). These findings demonstrate that obesity-driven NK cell impairment is reversible, highlighting metabolic programming as a tractable target to enhance antiviral immunity in high-risk populations.

## DISCUSSION

Despite annual IAV vaccine availability, IAV continues to cause significant morbidity and mortality worldwide, particularly among high-risk populations such as individuals with obesity. Obesity remains a major risk factor for severe IAV disease, yet the innate immune mechanisms underlying this vulnerability are unclear. Here, we identify NK cells as critical early antiviral effector cells whose protective function is abrogated in obesity. NK cells from DIO mice have severely blunted effector function responses, contributing to increased disease severity and loss of NK cell-mediated protection. Mechanistically, obesity rewires both the lung microenvironment and NK cell metabolic programming, resulting in lipid accumulation, mTOR dysfunction, and metabolic inflexibility. Importantly, these defects are reversible, as short-term weight loss restores NK cell effector function, metabolic signaling, and protection against IAV infection. Collectively, these findings reveal an immune-metabolic mechanism of NK cell dysfunction in obesity that contributes to increased IAV susceptibility and disease severity.

Previous studies have shown that obesity impairs NK cell effector functions in both human and murine systems, particularly in the context of cancer^20,59–61^. *Ex vivo* studies using NK cells isolated from obese individuals demonstrated defects in cytotoxicity, cytokine production, and metabolic pathways and these findings were corroborated with *in vivo* murine tumor models^59^. However, the extent to which obesity alters NK cell function during acute viral infection, particularly within the lung microenvironment, has remained poorly defined. Recent work demonstrated that obesity establishes an inflammatory and transcriptionally reprogrammed immune landscape in the lung and white adipose tissue during IAV infection, characterized by altered macrophage responses^62^. Collectively, prior work demonstrates that obesity drives NK cell dysfunction and lung immune reprogramming during IAV infection, yet the contribution of NK cells as early antiviral effectors and the mechanisms underlying their dysfunction in the obese lung microenvironment had not been defined.

Our findings demonstrate that obesity-driven NK cell dysfunction exhibits tissue specificity. While a prior study using a DIO model and a mouse-adapted H1N1 virus strain (A/Puerto Rico/ 08/1934; PR8) infection reported increased mortality and impaired NK cell cytotoxicity^63^, the mechanisms underlying this defect remained undefined. Our data expands upon these observations by revealing that the lung microenvironment shapes NK cell responses with distinct transcriptional and phenotypic remodeling of NK cells across developmental (bone marrow), systemic (spleen), and mucosal (lung) compartments. This is particularly relevant given that IAV infection is spatially heterogeneous and that antiviral control is determined locally at sites of viral replication. We identify delayed transcriptional induction of Tbet and defects in both cytotoxic and cytokine effector responses in lung NK cells in obesity and link these defects to alterations in the lung lipid microenvironment. Together, these findings supports a model in which tissue-local immune programming contributes to NK cell dysfunction during severe IAV infection in obesity.

In addition to antiviral functions, NK cells play a role in coordinating adaptive immunity through crosstalk with dendritic cells, macrophages, and T cells^14–17^. NK cell-derived IFN-γ promotes dendritic cell maturation and licenses optimal CD8⁺ T cell priming^16^. Therefore, early defects in NK cell activation and cytokine production in obese hosts may have broader consequences for immune coordination later in infection, contributing to impaired T cell immunity, delayed viral clearance, and dysregulated inflammation. These findings provide a potential mechanism for the well-described association between obesity and impaired vaccine responsiveness, in addition to increased susceptibility to severe respiratory viral infections. Given that NK cells serve as critical early responders to multiple respiratory pathogens^10,64^, our data suggest that obesity may establish a shared immune-metabolic state in the lung that broadly compromises early antiviral defense at mucosal sites and may impact recall responses to reinfection.

Cellular immune function is tightly coupled to membrane lipid composition, particularly the balance between the amount of saturated and unsaturated fatty acids in phospholipids^52,65^. We found that obesity and IAV infection increase a shift toward phospholipid species that contain a monounsaturated fatty acid in the lung. Phospholipid species including PC, PG, and PE, containing oleic (18:1) or palmitoleic (16:1) acyl chains regulate membrane fluidity, lipid raft organization, and can modulate immune signaling. In T cells, increased phospholipids containing 18:1 impacts TCR signaling, immune synapse engagement, and effector functions^66,67^. Lipid rafts are also essential for NK cell activation receptor organization, and a recent study found that PC 36:1 found in tumor ascites disrupts plasma membrane order and disrupts NK cell cytotoxicity and impacted lipid droplet formation^57^. This lipid remodeling is increasingly recognized as a mechanism that regulates immune responses and shapes lymphocyte effector functions by altering membrane organization, signaling complexes, and limiting metabolic fitness. We found increased PCs, PGs, and PEs containing monounsaturated fatty acids in the lung, however, the mechanism of how each of these changes contribute to NK cell-specific dysfunction, such as plasma versus mitochondrial membrane reorganization or impact on mitochondrial respiration are an area of current future research. Moreover, defining how these lipids change during infection, how they spatially align with sites of active IAV replication, and how they shape the function of other immune and non-immune cell populations within the lung will be critical for understanding how the lipid microenvironment impacts antiviral immunity.

It is well established that serum triglycerides are elevated in humans with obesity as well as in DIO obese mouse models, and we likewise observed significantly increased triglyceride abundance in obese mouse lungs by both bulk and spatial lipidomics. Triglycerides are broken down by lipases into free fatty acids by lipolysis, which can be taken up by lipid transporters such as CD36. We found that obesity drives excessive lipid uptake and storage in NK cells resulting in intracellular lipid accumulation that is associated with impaired mTOR signaling, mitochondrial dysfunction, and reduced metabolic flexibility, collectively limiting the energetic capacity required to sustain antiviral effector functions. Notably, prior studies have demonstrated that exposure to LCFAs is sufficient to suppress NK cell cytotoxicity and inhibit mTOR signaling, establishing intracellular lipid accumulation as a direct regulator of NK cell metabolic fitness^56,58,59^. Consistent with this model, NK cells from obese humans exhibit increased CD36 expression and elevated intracellular lipid accumulation, as assessed by LipidTOX and BODIPY staining, which correlates with impaired effector function^59^. Lipid droplets can be converted into energy through beta (β)-oxidation^68,69^. NK cells typically rely on β-oxidation during quiescence before switching to aerobic glycolysis as a primary energy source upon activation^70^. However, in obese mice we observed increased lipid droplet accumulation and reduced glycolytic activity in NK cells during IAV infection, raising the possibility that obese NK cells are metabolically stuck in a fatty acid-dependent state or have defects in β-oxidation that further suppresses mTOR activation and effector function. Together, these findings identify the lung lipid microenvironment as a key determinant of NK cell metabolic programming and antiviral responses during IAV infection.

We found that the lipid transporter CD36 is significantly elevated on obese NK cells, likely contributing to the increased lipid accumulation. Increased CD36 expression was previously found on peripheral human obese NK cells and linked to increased lipid accumulation and dampened effector functions^56,59^. However, NK cells express other lipid transporters, such as SCARB1 and several fatty acid binding proteins (FABP)^25,70^. Future research should investigate the contribution of different lipid transporters, including but not limited to CD36, on lipid accumulation and metabolic programming of NK cells during IAV infection in the lung. We also found that CD36 is highly expressed in obese lung tissue and may influence not only NK cells but also macrophages, epithelial cells, and endothelial cells, collectively reshaping pulmonary microenvironment. This could be important for the development of new therapeutics for IAV infection in high-risk populations.

A key translational implication of our study is that obesity-associated NK cell dysfunction is reversible. Short-term weight loss restored NK cell metabolic signaling, effector function, and protection to IAV infection, indicating that immune-metabolic remodeling is dynamic rather than permanent. These findings raise questions about how quickly immune function recovers following improvement of metabolic health and whether modest weight loss alone may be sufficient to restore antiviral immunity. As GLP-1 receptor agonists and other metabolic therapeutics become widely used, it will be important to understand how improvements in metabolic health translate into immune recovery. More broadly, our data suggests metabolic health is a modifiable determinant of antiviral immunity, with potential implications for vaccine responsiveness, pandemic preparedness, and infection-associated disease severity in people with obesity.

## Limitations of the study

This study has several limitations that should be considered. The majority of experiments were performed in male DIO mice, which consistently develop robust metabolic dysfunction and enabled reproducible modeling of obesity-associated immune impairment^28^. However, sex-specific differences in lipid metabolism and immune responses are well documented, and future studies in female mice will be important to determine whether similar immune-metabolic mechanisms constrain NK cell function across sexes. We used the pandemic H1N1 strain A/California/04/2009 to model a clinically relevant, non-lethal influenza infection, and while this approach allowed us to interrogate early antiviral immune responses under physiologic disease conditions, extending these findings to more severe infection models and additional influenza subtypes will be necessary to determine whether lipid-driven NK cell dysfunction similarly constrains immunity across a broader spectrum of disease severity. Although spatial metabolomics enabled resolution of lipid remodeling at sites of active viral replication, our analysis provides a snapshot during acute infection, and longitudinal studies will be required to define the temporal dynamics of lipid dysregulation across the course of infection. Moreover, while our data support a model in which lung lipid remodeling constrains NK cell metabolism and effector function, we have not yet directly established causality for individual lipid species or defined the cellular sources of excess lipids in infected lung regions. Finally, although our study is supported by previously published human data demonstrating increased CD36 expression and lipid accumulation in NK cells from obese individuals^59^, direct interrogation of lipid remodeling within the human lung during viral infection was not feasible. Future studies using human lung tissue platforms, including precision-cut lung slices and airway or alveolar organoid systems, will be essential to define how obesity shapes the pulmonary lipid microenvironment and programs NK cell function in a clinically relevant setting.

## METHODS

### RESOURCE AVAILABILITY

#### Lead Contact

Further information and requests for resources and reagents should be directed to and will be fulfilled by the Lead Contact, Dr. Stacey Schultz-Cherry.

#### Material Availability

Materials generated in this study are available from the lead contact upon request.

#### Data and Code Availability

Spatial lipidomics datasets will be made publicly available through Metaspace2020 (https://metaspace2020.org/) upon publication acceptance.

#### Experimental Model Details

##### Mice

All procedures were approved by the Institutional Biosafety Committee and the Animal Care and Use Committee at St. Jude Children’s Research Hospital in compliance with the Guide for the Care and Use of Laboratory Animals (protocol no. 513). These guidelines were developed by the Institute of Laboratory Animal Resources and approved by the Governing Board of the U.S. National Research Council. Mice were kept under 12-hour/12-hour light/dark cycles at an ambient temperature of 20°C and 45% humidity. They had continuous access to diet and water. When required, either for humane endpoints or time point collection, mice were euthanized following the American Veterinary Medical Association guidelines. Wildtype (WT) C57BL/6 male mice (Jackson Laboratory) were either placed on a standard diet (SD, Lab Diets 5001, 15% fat, 30% protein, 55% carbohydrate) or high-fat diet (HFD, Research Diets D12492, 60% fat, 20% protein, 20% carbohydrate) or diet-induced obesity (DIO) male mice in a WT C57BL/6 background were ordered directly from Jackson Laboratories. All mice were on their prospective diets for at least 12 weeks.

##### Cells and Viruses

MDCK cells (American Type Culture Collection, CCL-34) were cultured in Dulbecco’s minimum essential medium (DMEM; Corning) supplemented with 2 mM GlutaMAX (Gibco) and 10% fetal bovine serum (FBS; HyClone) and grown at 37°C with 5% CO_2_. A/California/04/2009 (H1N1) viruses were kind gifts from R. Webster at St. Jude Children’s Research Hospital. The virus was propagated in the allantoic cavity of 10-day-old specific pathogen-free embryonated chicken eggs at 37°C. Briefly, allantoic fluid was cleared by centrifugation following harvest and then stored at −80°C. Viral titers were determined by TCID_50_ analysis.

##### Viral Titer Determination

Viral titers were quantified by TCID_50_ assays as previously described. Briefly, confluent MDCK cells were infected in duplicate or triplicate with 10-fold serial dilutions of virus stocks or tissue homogenates (bead beat in 500 μl of PBS) in 100 μl of minimal essential medium (MEM) plus 0.75% bovine serum albumin (BSA) and tosylsulfonyl phenylalanyl chloromethyl ketone (1 μg/ml)–treated trypsin (H1N1 samples only). After 3 days of incubation at 37°C and 5% CO_2_, 50 μl of the supernatant was combined and mixed with 50 μl of 0.5% packed turkey red blood cells diluted in PBS for 45 min at room temperature and scored by HA endpoint. Infectious viral titers were calculated using the Reed-Muench method^71^.

##### Mouse Viral Infectivity

Mice were challenged by intranasal inoculation with 1000 TCID_50_ of A/California/04/2009 diluted in PBS to a total volume of 25 μl that were lightly anesthetized with inhaled isoflurane or with 25 μl of PBS as a mock control. Body weight, clinical scores, and survival were monitored daily for 14 days. Moribund mice that lost more than 30% body weight and/or reached clinical scores of greater than 4 were humanely euthanized. Clinical signs were scored as follows: 0, no observable signs; 1, active, squinting, mild hunch, or scruffy appearance; 2, squinting and hunching, ruffled fur, but active; 3, excessive hunching, squinting, visible weight loss, not active when stimulated; 4, not active when stimulated, sunken face and severe weight loss, shivering, rapid breathing, and moribund; and 5, death.

##### Flow Cytometry and Cell Sorting

Mouse spleens, lungs, and bone marrow were harvested and processed into single-cell suspensions for flow cytometric analysis.

###### Spleen

Spleens were harvested, washed in PBS, and transferred to complete RPMI medium. Tissues were mechanically dissociated through a 70-μm cell strainer using the plunger end of a 3–5 mL syringe. The strainer was washed with 5 mL PBS, and the cell suspension was centrifuged at 500 × g for 10 min at 4°C. Pellets were resuspended in 3 mL of 1× red blood cell (RBC) lysis buffer (BioLegend) for 2 min. Lysis was quenched with 3 mL PBS, followed by centrifugation at 500 × g for 5 min at 4°C. Cells were resuspended in 10 mL complete RPMI medium to generate single-cell splenocyte suspensions.

###### Bone marrow

Femurs and tibias were resected, cleared of surrounding tissue, and severed at the proximal and distal ends to expose the medullary cavity. Bone marrow was flushed with cold PBS using a 25G needle and syringe. The resulting cell suspension was mechanically dissociated through a 70-μm cell strainer and subsequently filtered through a 100-μm mesh filter to remove debris. Cells were washed with PBS and subjected to RBC lysis using 1× RBC Lysis Buffer (BioLegend) for 4 min at room temperature. Lysis was quenched with excess PBS, and cells were centrifuged twice at 350 × g for 10 min. Pellets were resuspended in PBS, and viable cells were counted using a hemocytometer.

###### Lung

Lungs were harvested aseptically and minced into small fragments using sterile scissors. Tissue was digested in 1 mL digestion buffer consisting of RPMI supplemented with 400 U/mL type I collagenase (Worthington, LS004196) and 50 U/mL DNase I (Worthington, LS0002145) at 37°C with agitation (220 rpm) for 40 min. Digestion was quenched by addition of fetal bovine serum (FBS) to a final concentration of 10%. Digested tissue was passed through a 100-μm cell strainer and mechanically dissociated using the plunger end of a 5 mL syringe. The strainer was washed with 5 mL PBS, and the filtrate was centrifuged at 300 × g for 5 min at 4°C. Pellets were resuspended in 1 mL RBC lysis buffer (BioLegend; 1:10 dilution in water) for 2 min at room temperature, quenched with 2 mL PBS, and centrifuged at 300 × g for 5 min at 4°C. Cells were resuspended in 5 mL RPMI supplemented with FBS, and viable cells were counted.

###### Flow Staining

Antibodies used in this study, including clone, fluorophore, and vendor information, are listed in the **Key Resources Table**. Cells were permeabilized using the Foxp3 Fixation/Permeabilization Kit (eBioscience) for transcription factor staining and phosphorylation flow staining or Cytofix/Cytoperm (BD Biosciences) for cytokine staining and all other staining protocols. To assess NK cell cytokine production and activation, approximately 2.5 × 10^6^ cells/sample were stimulated with either 1 μg/mL leukocyte activation cocktail with GolgiPlug (BD Bioscience) or with eBioscience™ Cell Stimulation Cocktail (500X) at 37°C for 4 hours and stained intracellular markers following fixation and permeabilization. LipidTOX staining was conducted according to the manufacturer’s instructions. Flow cytometry was conducted using either a Fortessa (BD Biosciences) or Aurora (Cytek) flow cytometer. Flow cytometry data were analyzed using FlowJo software (Treestar), and flow cytometry antibody panels were designed using Fluorofinder.

#### Immune cell depletion

For *in vivo* NK cell depletion, mice were administered 300 μg anti-NK1.1 monoclonal antibody (PK136, BioXCell) or the corresponding InVivoPlus mouse IgG2a isotype control (BioXCell) by intraperitoneal injection -2, -1, day of infection, +1, then every other day until the conclusion of the experiment. Depletion efficiency was assessed by flow cytometric analysis of single-cell suspensions prepared from spleen and lungs at the time of infection using staining for the NK cell markers NK1.1 and DX5. Validation of NK cell depletion is shown in Supplemental Figure 1.

#### Seahorse metabolic flux analysis of lung NK cells

Lung single-cell suspensions were prepared as described above and processed immediately for NK cell isolation. Dead cells were removed using the Mouse Dead Cell Removal Kit (BioLegend) according to the manufacturer’s instructions. NK cells were subsequently enriched by two sequential rounds of negative selection using a magnetic bead-based Mouse NK Cell Isolation Kit (BioLegend) following manufacturer’s instructions. Purified NK cells were washed, resuspended in PBS and counted using trypan blue using a hemacytometer. For metabolic flux analysis, NK cells were resuspended in Seahorse XF RPMI assay medium (pH 7.4) supplemented with glucose (10 mM), sodium pyruvate (1 mM), and glutamine (2 mM) (Agilent 103681-100). Cells were either plated at 50,000 cells per well or normalized to cell count thereafter in duplicate in a poly-D-lysine–coated Seahorse XF HS PDL Miniplate and centrifuged at 200 × g for 5 min to promote adherence. Oxygen consumption rate (OCR) and extracellular acidification rate (ECAR) were measured using a Seahorse XF Mini and XF T Cell Metabolic Profiling Kit (Agilent 103772-100) according to the manufacturer’s instructions. Data were quality controlled and analyzed using Microsoft Excel and Agilent Seahorse Analytics software.

#### Lipid mass spectrometry

Whole lung homogenates were used to extract lipids using the Bligh and Dyer method^72^. Lipid extracts were resuspended in chloroform/methanol (1:1). For the free fatty acid analysis, the sample was spiked with U-^13^C_18_-oleic acid (Cambridge Isotope laboratories, Inc.) as an internal standard. Lipid species were analyzed using a Shimadzu Prominence UFLC attached to a QTrap 4500 equipped with a Turbo V ion source (Sciex). Samples were injected onto an Acquity UPLC BEH HILIC, 1.7 μm, 2.1 × 150 mm column (Waters) at 45 °C with a flow rate of 0.2 ml/min. Solvent A was acetonitrile, and solvent B was 15 mM ammonium formate, pH 3.0. The HPLC program was the following: starting solvent mixture of 96% A/4% B; 0 to 2 min, isocratic with 4% B; 2 to 20 min, linear gradient to 80% B; 20 to 23 min, isocratic with 80% B; 23 to 25 min, linear gradient to 4% B; and 25 to 30 min, isocratic with 4% B. The QTrap 4500 was operated in the Q1 negative mode for free fatty acids and PG and in the positive mode for PC and PE. The negative ion source parameters for Q1 were as follows: ion spray voltage, −4500 V; curtain gas, 25 psi; temperature, 350 °C; ion source gas 1, 40 psi; ion source gas 2, 60 psi; and declustering potential, −40 V. The positive ion source parameters for Q1 were as follows: ion spray voltage, 5500 V; curtain gas, 25 psi; temperature, 350 °C; ion source gas 1, 40 psi; ion source gas 2, 60 psi; and declustering potential, 40 V. The system was controlled by the Analyst software (Sciex). The sum of the areas under each peak in the mass spectra was calculated, and the percentage of each molecular species present was calculated with LipidView software (Sciex). For free fatty acids levels, the data was compared to the internal standard for relative abundance.

The QTrap 4500 was used to perform product scans to verify the fatty acids present in a particular PC, PE, and PG molecular species. The ion source parameters for negative mode product scan were: ion spray voltage, -4500 V; curtain gas, 10 psi; collision gas, medium; temperature, 270 □C; ion source gas 1, 10 psi; ion source gas 2, 15 psi; declustering potential, - 40 V; and collision energy, -50 V.

#### MALDI-MSI Spatial Lipidomics

##### Tissue Preparation

Mice were perfused transcardially with 4% paraformaldehyde in PBS. Lungs were immediately inflated with 3.75% (hydroxylpropyl)methyl cellulose and 1.25% polyvinylpyrrolidone (HPMC-PVP) in PBS and snap frozen within a matrix comprised of 7.5% HPMC and 2.5% PVP using LN2-cooled 2-methylbutane. Tissues were stored at -80 degrees until cryosectioned at -20 degrees. 10μm thick sections were thaw mounted onto indium tin oxide (ITO) slides for MALDI analyses and stored at -80 until use.

##### Sample Preparation for MALDI MSI Analysis

ITO slides with lung sections were brought to room temperature in a lyophilizer for 15 minutes prior to application of MALDI matrix. A solution of 40 mg/mL 2,5-dihydroxybenzoic acid (DHB) with 500μM of ammonium fluoride was dissolved in a 70% methanol solution. MALDI matrix was applied using an HTX M5 sprayer (HTX Technologies). Briefly, matrix was sprayed homogenously across slides over 9 passes with a 20 sec drying time between passes (nozzle temperature of 75°C and bed temperature of 60°C, a solvent flow rate of 0.2 mL/min and velocity set to 1920 mm/min, a gas flow rate of 3L/min at 10 psi with a track spacing of 2 mm and spray pattern of CC; nozzle height set to 40 mm). After matrix application, samples were immediately analyzed.

##### MALDI MSI Analysis

MALDI MSI analyses were performed using a Bruker timsTOF fleX mass spectrometer (Bruker Daltonics). Before acquisition, external calibration was completed using Agilent ESI tune mix for ESI-TOF. Transfer settings were a MALDI plate offset of 50 V, deflection 1 delta of 70 V, 350 V peak-to-peak (Vpp; funnel 1 RF), 350 Vpp (funnel 2 RF), and 350 Vpp (multipole RF). Focus pretime-of-flight (TOF) transfer time was set at 70 μs and prepulse storage at 10 μs. The quadrupole ion energy was 10 eV with a low mass of m/z 100 Da. Collision cell energy was 10 eV with the collision RF set to 1500 Vpp. Data was acquired in positive ionization mode over a range of 100 -1800 Da using a Smartbeam 3D laser firing 500 laser shots per pixel at a frequency of 10,000 Hz. A laser raster size of 20 μm operating in beamscan mode at a laser power of 90% with 0% power boost was utilized. Ion images were acquired with a spatial raster of 20 μm. After acquisition, data was processed using Bruker SCiLS Lab (version 2026A) and normalized to the total ion chromatogram (TIC). SCiLS lab was utilized to convert data files to.imzML and.ibd using no normalization and the top 1,500 features for tentative identifications of lipids using Metaspace2020^73^.

#### Immunofluorescence and confocal microscopy

##### Tissue Immunofluorescence

Lung tissues were cryosectioned at 10μm thickness and allowed to air dry for 30min. Sections were washed in PBS for 5min prior to incubation in blocking buffer comprised of PBS containing 1% bovine serum albumin. Antibody labeling was performed overnight in PBS containing 1% BSA using the following antibodies: goat anti-NP (US Biological; catalog 17650-05E), rat anti-NKp46 (Biolegend; clone 29A1.4, catalog 137601), rat anti-CD36 (Biolegend; clone ZND36-6, catalog 163002). Sections were washed in PBS prior to detection with species-specific secondaries followed by mounting in Vectashield Vibrance mounting medium (Vector labs). Widefield fluorescence imaging was performed using a Ti2 inverted microscope (Nikon Instruments) equipped with an Orca Fusion sCMOS camera (Hamamatsu), SOLA II LED lightsource (Lumencorp), and excitation/emission filters as appropriate. Images were acquired and analyzed using NIS Elements software, version 5.42.06 (Nikon Instruments)

##### Cell Immunofluorescence

Purified cells were allowed to settle onto poly-L-Lysine coated coverslips (EMSdiasum) in complete media at 37 degrees for 15 min prior to fixation with 4% paraformaldehyde and 0.1% glutaraldehyde in PBS for 10min at RT. Cells were washed in PBS and blocked with PBS containing 1% BSA prior to incubation with rat anti-NKp46 (Biolegend) followed by detection with AF488-labelled rat-specific secondary antibody. Cells were imaged in PBS containing HCS LipidTox deep red reagent (ThermoFisher; catalog H34477) following the manufacturer’s suggested protocol. Cells were imaged using a Marianas spinning disk confocal (Intelligent Imaging Innovations) comprised of an inverted ObserverZ.1 microscope (Zeiss), CSU-W SoRa spinning disk (Yokogawa), Prime95B sCMOS camera (Photometrics) and 100x oil objective. 3D imaging was performed using a 2.8X SoRa mag changer, yielding 0.039μm/px. Images were subsequently deconvolved using a nearest-neighbors algorithm and Slidebook software, version 2025.2 (Intelligent Imaging Innovations).

### Statistical Analyses

Experiments were conducted in duplicate or triplicate and repeated at least twice. Representative results of single experiments are shown in the figures as indicated. All data visualization and statistical analyses were performed using GraphPad Prism v10.0. Sample sizes are indicated in the corresponding figure legends. Statistical significance was assessed using one-way ANOVA with Tukey’s post hoc test, two-way mixed-effects ANOVA with multiple-comparisons testing, or log-rank (Mantel-Cox) analysis for survival curves, as specified for each experiment.

### Data and materials availability

All data needed to evaluate the conclusions in the paper are present in the paper and/or the Supplementary Materials.

## KEY RESOURCE TABLE

| REAGENT or RESOURCE | SOURCE | IDENTIFIER |
| --- | --- | --- |
| **Antibodies** |  |  |
| Anti-mouse CD45.2<br>PerCP/Cyanine5.5 Clone: 104 | Biolegend | 109828 |
| Anti-mouse CD45.2<br>APC-Cyanine7 Clone: 104 | Biolegend | 109824 |
| Anti-mouse CD45<br>PE-Cyanine7 Clone: 30-F11 | Biolegend | 103113 |
| Anti-mouse NK1.1<br>PE-Cyanine5 Clone: PK136 | Biolegend | 108716 |
| Anti-mouse NK1.1<br>PE Clone: PK136 | Biolegend | 108708 |
| Anti-mouse C49b<br>APC Clone: DX5 | Biolegend | 108910 |
| Anti-mouse TCR beta chain<br>Brilliant Violet 421 Clone: H57-597 | Biolegend | 109229 |
| Anti-mouse CD36<br>BB515 Clone: CRF D-2712 | BD Biosciences | 565094 |
| Anti-human/mouse phospho-mTOR (Ser2448)<br>PE-Cyanine7 Clone: MRRBY | Invitrogen | 25-9718-42 |
| Anti-human/mouse phospho-AKT1 (Ser473)<br>PE Clone: SDRNR | Invitrogen | 11-9715-42 |
| Anti-mouse P-S6 RiboProt (S235/236) Alexa 488<br>Clone:D57.2.2E | Cell Signaling | 4803S |
| Anti-mouse CD98 (4F2)<br>PE/Cyanine 7 Clone: RL388 | Biolegend | 128214 |
| Anti-mouse CD69<br>APC/Cyanine7 Clone: H1.2F3 | Biolegend | 104525 |
| Anti-mouse/rat/human CD27<br>Brilliant Violet 605 Clone: LG.3A10 | Biolegend | 124249 |
| anti-mouse/human CD11b<br>Antibody PE/Cyanine7 Clone: M1/70 | Biolegend | 101215 |
| Anti-mouse/rat/human Glut-1<br>ALEXA FLUOR™ 647 Clone: 07-1401-AF647 | Millipore | 07-1401-AF647 |
| Anti-Tbet FITC Clone: 4B10 | Biolegend | 644811 |
| Anti-mouse Interferon Gamma<br>PE Clone: XMG1.2 | Biolegend | 505807 |
| Anti-mouse/human Granzyme B<br>Pacific Blue Clone: GB11 | Biolegend | 515408 |
| Anti-mouse/rat/human c-myc FITC Clone: SH1-26E7.1.3 | Miltenyi Biotec | 130-116-485 |
| Anti-mouse/human Ki-67 FITC Clone: 11F6 | Biolegend | 151212 |
| Anti-GATA3 mouse PE-CF594 Clone: :50-823 | BD Biosciences | 563510 |
| Anti-mouse EOMES PE Clone: W17001A | Biolegend | 157705 |
| FITC Rat IgG2a Isotype Control Clone: G013C12 | Biolegend | 402307 |
| PE Rat IgG1 Isotype Control Clone: G0114F7 | Biolegend | 401905 |
| Rat IgG2a kappa Isotype Control (eBR2a), PE, | eBioscience™ | 12-4321-80 |
| InVivoMAb anti-mouse NK1.1 | BioxCell | BE0036 |
| InVivoMAb mouse IgG2a isotype control | BioxCell | BE0085 |
| rat anti-NKp46 clone 29A1.4 | Biolegend | 137601 |
| goat anti-NP influenza protein | US Biological | 17650-05E |
| rat anti-CD36 clone ZND36-6 | Biolegend | 163002 |
| HCS LipidTox deep red reagent | ThermoFisher | H34477 |
| **Chemicals, peptides, and recombinant proteins** |  |  |
| Zombie Aqua™ Fixable Viability Kit | Biolegend | 423102 |
| UltraComp eBeads™ Compensation Beads | ThermoFisher (Invitrogen) | 01-2222-42 |
| Leukocyte Activation Cocktail, with BD GolgiPlug™ | BD Biosciences | 550583 |
| Purified Rat Anti-Mouse CD16/CD32 (Mouse BD Fc Block™) | BD Biosciences | 553142 |
| Trypan Blue Solution, 0.4% | ThermoFisher Scientific | 15250061 |
| DNase I | Worthington | LS002145 |
| Collagenase type I | Worthington | LS004196 |
| DIO Rodent Purified Diet w/60%Kcal Fat- Blue (58Y1) |  | 58126 |
| BD BD GolgiStop™ Protein Transport Inhibitor (containing Monensin) | Fisher Scientific | BDB554724 |
| eBioscience™ Cell Stimulation Cocktail (500X) | ebioscience | 00-4970-03 |
| RBC Lysis Buffer (10X) | Biolegend | 420301 |
| MojoSort™ Buffer (5X) | Biolegend | 480017 |
| **Critical commercial assays** |  |  |
| BD Cytotfix/cytoperm | BD Biosciences | 554714 |
| Transcription Factor Buffer Set | BD Biosciences | 562574 |
| MojoSort™ Mouse NK Cell Isolation Kit | Biolegend | 480049 |
| MojoSort™ Mouse Dead Cell Removal Kit | Biolegend | 480157 |
| HCS LipidTOX™ Green Neutral Lipid Stain | Thermo Fisher Scientific | H34475 |
| eBioscience™ Foxp3 / Transcription Factor Staining Buffer Set | eBioscience | 00-5523-00 |
| Triglyceride Colorimetric Assay Kit | Cayman Chemical | 10010303 |
| Seahorse XF RPMI assay medium pack | Agilent | 103681-100 |
| Seahorse XF HS Mini FluxPak (PDL plates) | Agilent | 103724-100 |
| Seahorse XF T Cell Metabolic Profiling Kit | Agilent | 103771-100 |
| **Experimental models: Organisms/strains** |  |  |
| C57BL/6J | Jackson Laboratory | Cat# 000664 |
| C57BL/6J DIO | Jackson Laboratory | Cat#380050 |
| A/California/04/2009 | Provided by R. Webster | N/A |
| **Software and algorithms** |  |  |
| Graphpad Prism 10 | Graphpad Prism Software |  |
| FlowJo v. 10.1 | Tree Star |  |
| METASPACE2020 | METASPACE2020 | https://metaspace2020.org |
| QuPath 0.5.1 | QuPath |  |
| BioRender | Biorender Company |  |
| Agilent Seahorse Analytics | Agilent |  |
| SCiLS Lab (version 2026A) | Bruker |  |

## SUPPLEMENTAL INFORMATION

Supplemental Figures 1-5.

## ACKNOWLEDGEMENTS

We are grateful to members of the Schultz-Cherry lab for many useful discussions. We are grateful to Tyler Ripperger for helpful discussions and feedback on data presentation, Maria Smith for assistance with technical support and early discussions that informed this work, and Elaine Tuomanen for careful review of the manuscript. We would like to thank the Animal Resource Center that assisted with blood chemistries, the Veterinary Pathology Core that assisted with animal handling, and the Flow Cytometry Core for guidance and expertise. This work was supported by the National Institute of Allergy and Infectious Diseases, National Institutes of Health, Department of Health and Human Services, under contract no. 75N93021C00016 (S.S.-C.); the American Lebanese Syrian Associated Charities (ALSAC) (S.S.-C.); the National Institute of Health, Institutional Postdoctoral Training Grants (T32) Infectious Disease Therapeutics T32AI106700-08 (P.H.B.); and the National Institute of Health, Ruth L. Kirschstein Postdoctoral Individual National Research Service Award F32AI183804 (P.H.B.).

## AUTHOR CONTRIBUTIONS

Conceptualization: P.H.B, S.S.-C.; Methodology: P.H.B, M.F., C.G., A.M., E.S., J.B. S.S.-C.; Formal Analysis: P.H.B, M.F., E.S.; Investigation: P.H.B, M.F., L.R., T.B., C.G., B.L., A.M, E.S. S.S.-C.; Project Administration: P.H.B, S.S.-C.; Supervision: P.H.B, S.S.-C.; Writing – Original Draft: P.H.B. Writing – Revision and Edits: P.H.B., S.S.-C., M. F., A.M. ; Funding Acquisition: P.H.B., S.S.-C.

---

**Supplementary Figure 1.**
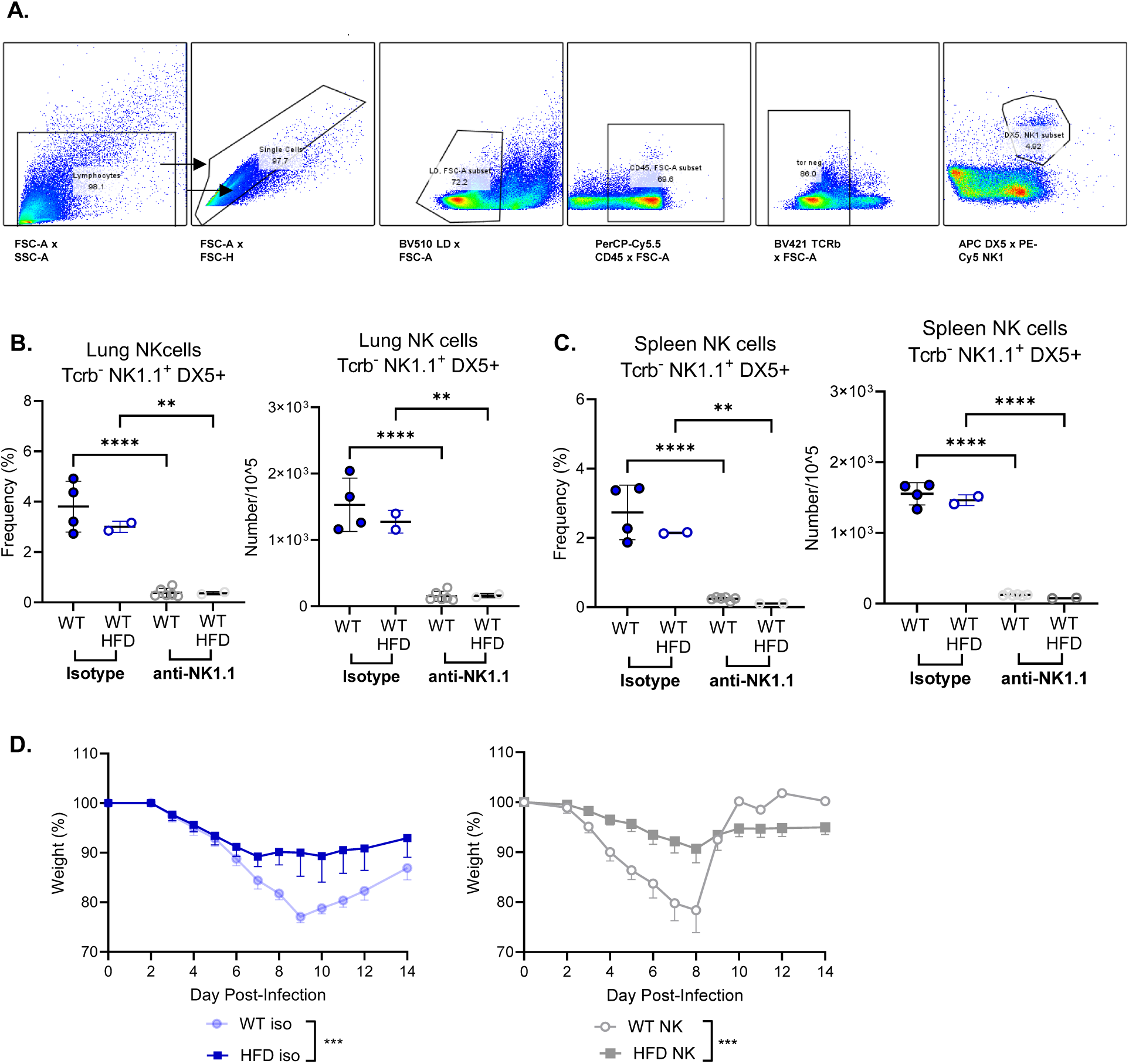
NK depletion validation and impact of NK cell depletion on weight change and following influenza infection. WT mice were placed on standard diet (SD) or 60% high-fat diet (HFD) for a minimum of 12 weeks. SD and HFD mice were inoculated by intranasal (IN) infection with 1000 tissue culture infectious dose-50 (TCID_50_) of A/California/04/2009 (CA/09 H1N1) virus or PBS as a mock control. **(A-C)** NK cells were depleted with 300 μg of anti-NK1.1 PK136 antibody or an IgG2a isotype control days -2, -1, day of infection (d0) and then lungs or spleens were resected and processed for flow cytometry. **(A)** Example gating of lung NK cells **(B)** Frequency and number of NK cells (CD45^+^ Tcrβ^-^ NK1.1^+^ DX5^+^) assessed by flow cytometry in the lung and **(C)** in the spleen. **(D)** NK cells were depleted with 300 μg of anti-NK1.1 PK136 antibody or an IgG2a isotype control days -2, -1, day of infection (d0), +1, and then every other day throughout the duration of the experiment and weight loss over time was assessed. Data are shown as mean ± SD **(B, C)** or SEM **(D)**. Statistical significance was calculated using a One-way ANOVA with multiple comparison’s test **(B, C)** and a Two-way ANOVA **(D)**, ** p < .01; *** p < .001; **** p < .0001; ns = not significant.

**Supplementary Figure 2.**
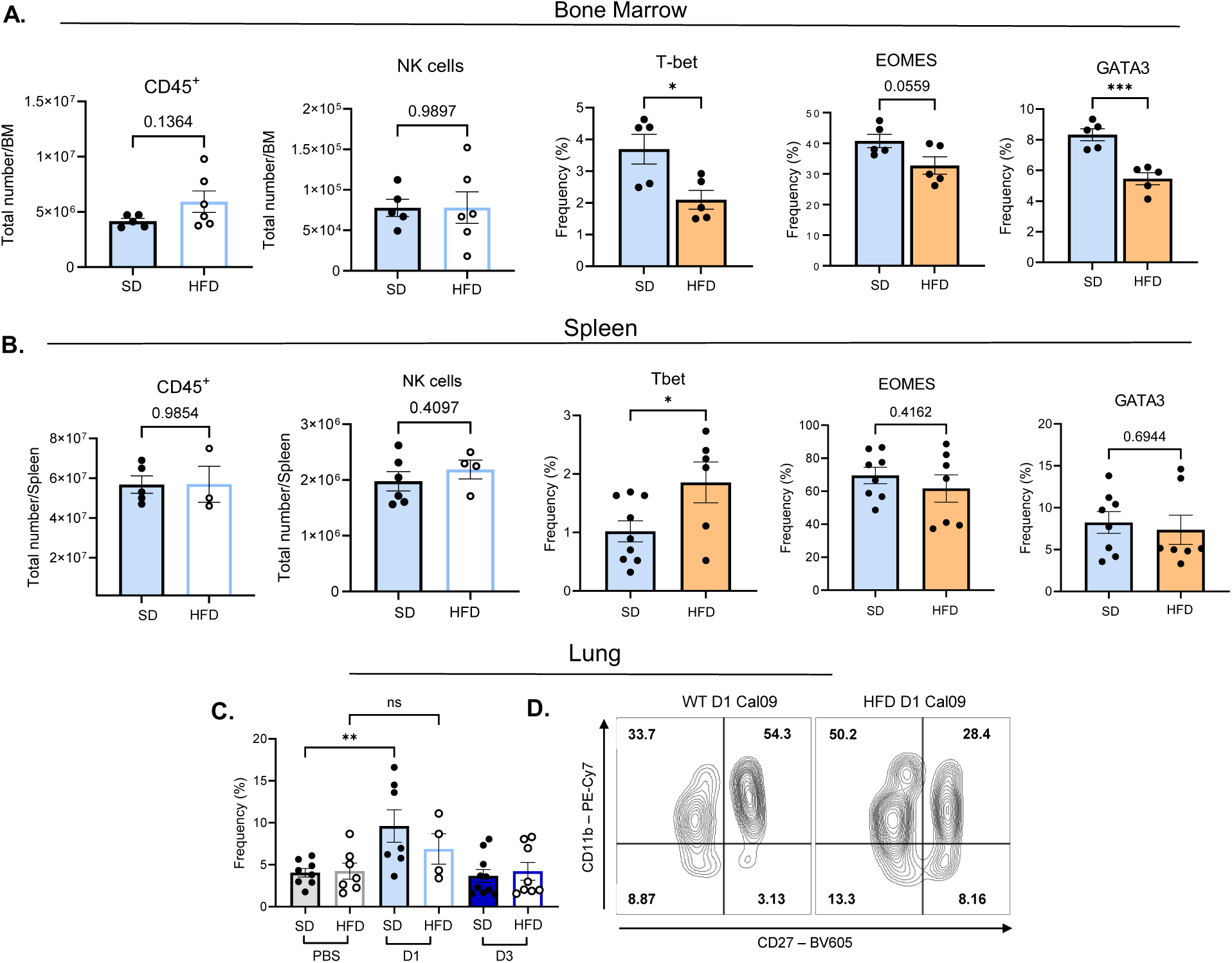
Diet-induced obesity alters NK cell maturation and transcription factor expression at baseline differently based on immune compartment. WT mice were placed on SD or 60% HFD for a minimum of 12 weeks. **(A)** Bone marrow was resected and processed for flow cytometry, including the total number of immune cells (CD45^+^), NK cells (CD45^+^ Tcrβ^-^ NK1.1^+^) and intracellular staining for NK cell transcription factors including Tbet, EOMES, and GATA3, or **(B)** assessed in the spleen. **(C, D)** SD and HFD mice were inoculated by IN infection with 1000 TCID_50_ of CA/09 H1N1 virus or PBS as a mock control and assessed for **(C)** frequency of lung NK cells. **(D)** Representative contour plot of CD11b and CD27 gating on lung NK cells. Data are shown as mean ± SEM **(D)**. Statistical significance was calculated using a student’s t test with Welch’s correction **(A, B)** or a One-way ANOVA with multiple comparison’s test **(C)** * p < .05; ** p < .01; *** p < .001.

**Supplementary Figure 3.**
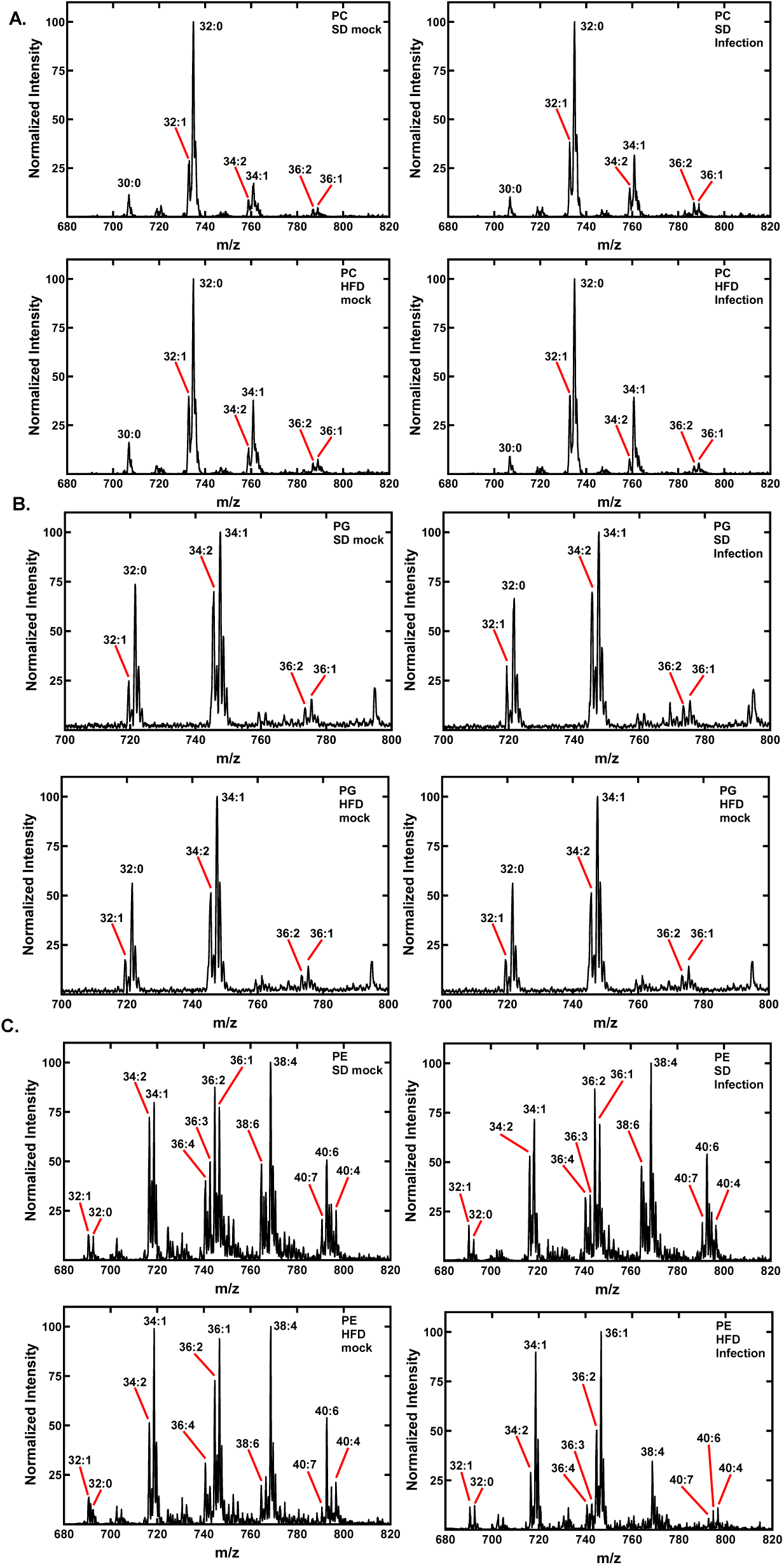
Representative mass spectrometry plots of phospholipid profiles in SD or HFD mice with and without influenza infection. WT mice were placed on SD or 60% HFD for a minimum of 12 weeks. SD and HFD mice were inoculated by IN infection with 1000 TCID_50_ of CA/09 H1N1 virus or PBS as a mock control. Representative mass spectrometry lipidomics plots with normalized intensity of **(A)** Phosphatidylcholine (PC) **(B)** Phosphatidylglycerol (PG) or **(C)** phosphatidylethanolamine (PE).

**Supplementary Figure 4.**
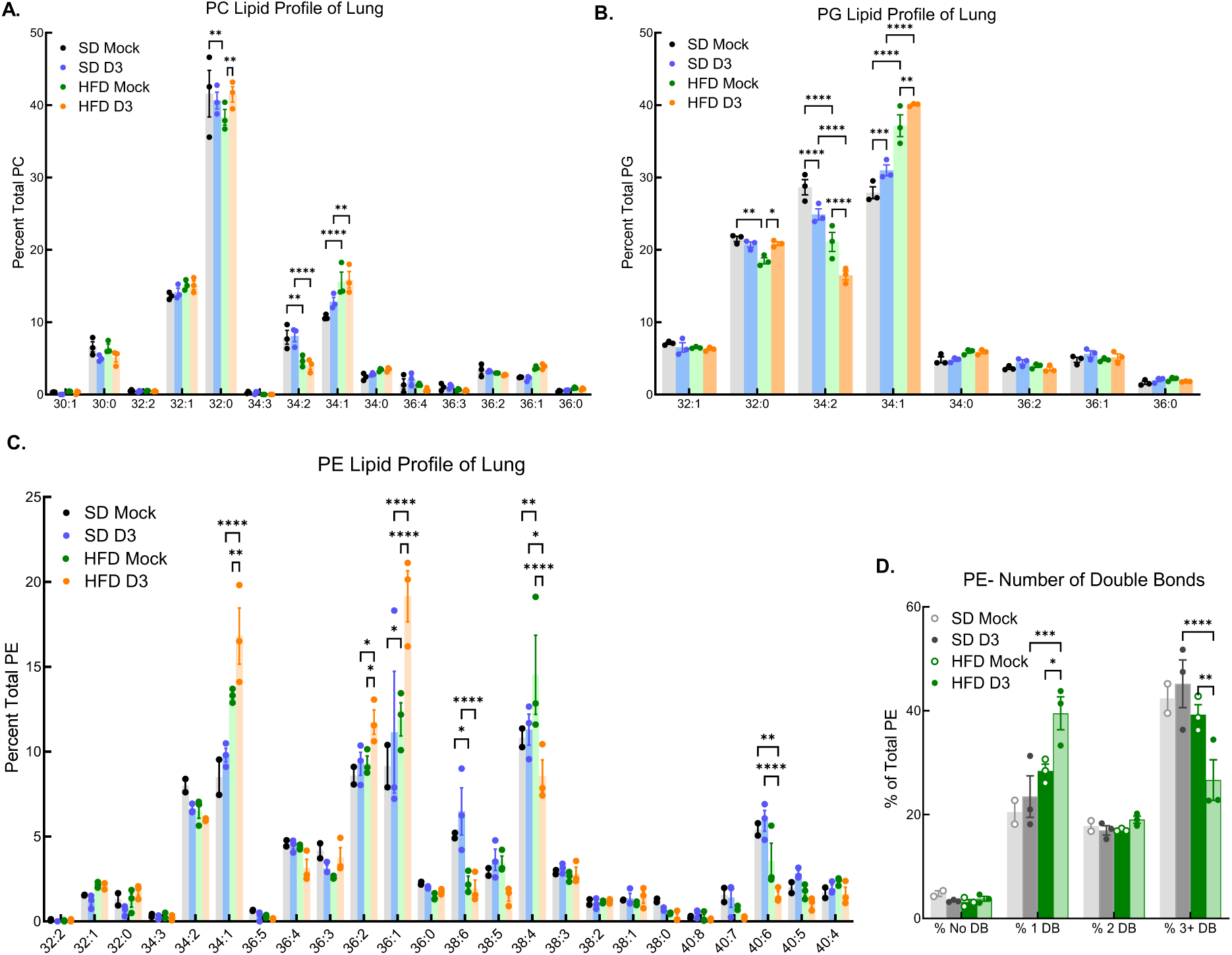
Relative abundance of phospholipid lung lipid profiles in SD or HFD mice with and without IAV infection. WT mice were placed on SD or 60% HFD for a minimum of 12 weeks and were intranasally inoculated with 1000 TCID_50_ of CA/09 H1N1 virus or PBS (mock control). At 3 dpi, lungs were resected and processed for bulk lipidomics. **(A)** Relative abundance of PC **(B)** PG **(C)** PE and **(D)** Relative percentage of number of double bond distribution among all analyzed PE. Data are shown as mean ± SEM. Statistical significance was calculated using a two-way ANOVA with multiple comparison’s **(A-D)**. * p < .05; ** p < .01; *** p < .001; **** p < .0001.

**Supplementary Figure 5.**
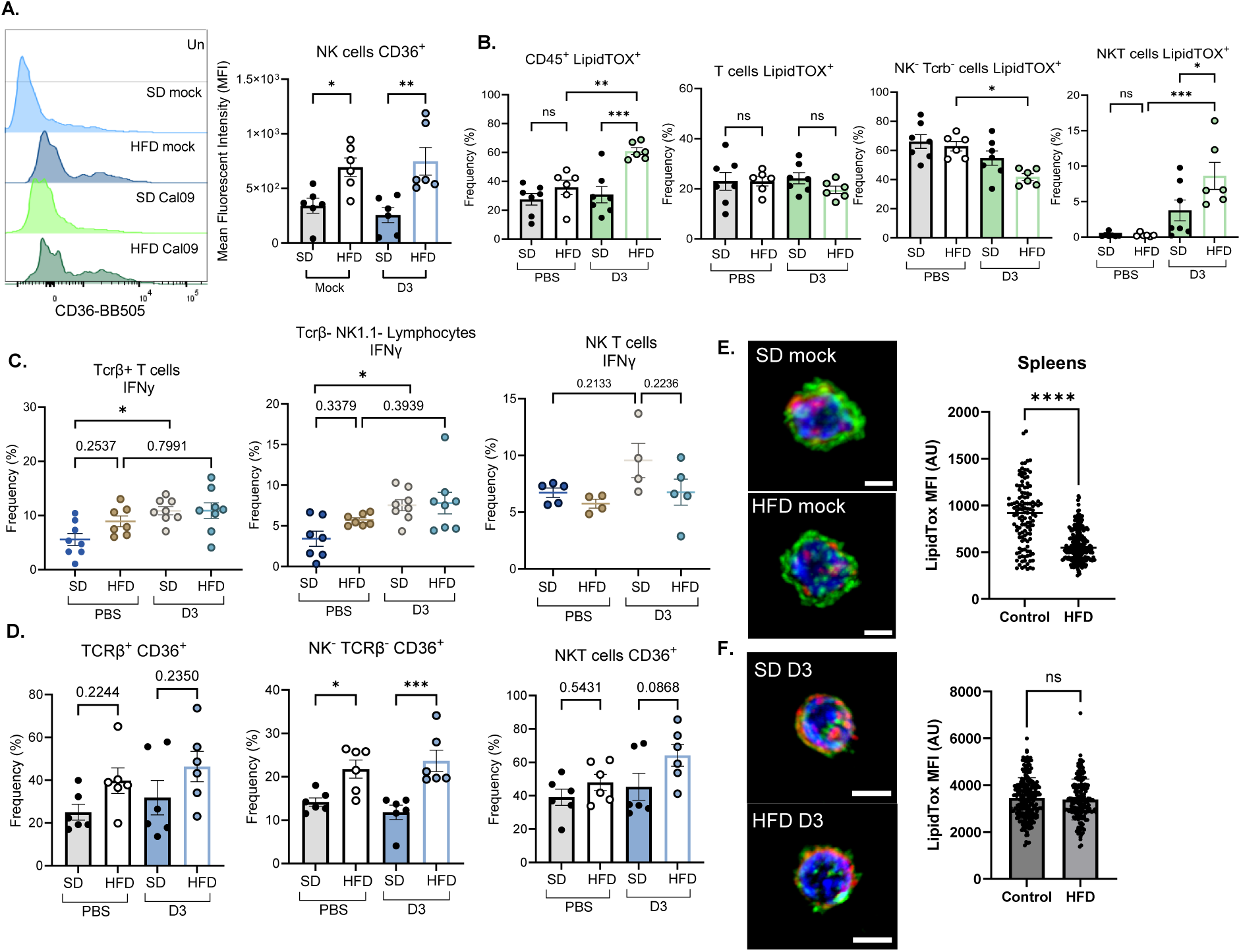
Lipid and immune signatures across multiple immune cell types. WT mice were placed on SD or 60% HFD for a minimum of 12 weeks and were intranasally inoculated with 1000 TCID_50_ of CA/09 H1N1 virus or PBS (mock control). At 3 dpi, lungs were resected and processed for flow cytometry. **(A)** Representative MFI plot and MFI of CD36 expression on NK cells **(B)** LipidTOX staining for total immune cells, (CD45^+^), T cells (CD45^+^ Tcrβ^+^ NK1.1^-^), other immune cells (CD45^+^ Tcrβ^-^ NK1.1^-^), and NK T cells (CD45^+^ Tcrβ^+^ NK1.1^+^) **(C)** IFN-γ expression by T cells (CD45^+^ Tcrβ^+^ NK1.1^-^), other immune cells (CD45^+^ Tcrβ^-^ NK1.1^-^), and NK T cells (CD45^+^ Tcrβ^+^ NK1.1^+^) and **(D)** CD36 expression. **(E, F)** At 3 dpi with PBS (mock) **(E)** or IAV **(F)**, spleens were resected and processed for single cell sorting to isolate lung NK cells by magnetic activated cell sorting for IF imaging of NK cells (NKp46+, green), DAPI (blue) and LipidTOX (red) with representative images and MFI quantitation of LipidTOX staining. Data are shown as mean ± SEM. Statistical significance was calculated using a One-way ANOVA with Tukey’s multiple comparison’s **(A-D)** or a student’s t test **(E, F)**. * p < .05; ** p < .01; *** p < .001; ns = not significant.

## REFERENCES

1. Krammer, F., Smith, G.J.D., Fouchier, R.A.M., Peiris, M., Kedzierska, K., Doherty, P.C., Palese, P., Shaw, M.L., Treanor, J., Webster, R.G., et al. (2018). Influenza. Nat Rev Dis Primers 4, 3. 10.1038/s41572-018-0002-y.

2. Influenza (seasonal) https://www.who.int/news-room/fact-sheets/detail/influenza-(seasonal).

3. Smith, M., Honce, R., and Schultz-Cherry, S. (2020). Metabolic Syndrome and Viral Pathogenesis: Lessons from Influenza and Coronaviruses. Journal of Virology 94, 10.1128/jvi.00665-20. 10.1128/jvi.00665-20.

4. Honce, R., and Schultz-Cherry, S. (2019). Impact of Obesity on Influenza A Virus Pathogenesis, Immune Response, and Evolution. Front Immunol 10, 1071. 10.3389/fimmu.2019.01071.

5. Louie, J.K., Acosta, M., Samuel, M.C., Schechter, R., Vugia, D.J., Harriman, K., Matyas, B.T., and the California Pandemic (H1N1) Working Group (2011). A Novel Risk Factor for a Novel Virus: Obesity and 2009 Pandemic Influenza A (H1N1). Clin Infect Dis 52, 301–312. 10.1093/cid/ciq152.

6. Kerkhove, M.D.V., Vandemaele, K.A.H., Shinde, V., Jaramillo-Gutierrez, G., Koukounari, A., Donnelly, C.A., Carlino, L.O., Owen, R., Paterson, B., Pelletier, L., et al. (2011). Risk Factors for Severe Outcomes following 2009 Influenza A (H1N1) Infection: A Global Pooled Analysis. PLOS Medicine 8, e1001053. 10.1371/journal.pmed.1001053.

7. Honce, R., Karlsson, E.A., Wohlgemuth, N., Estrada, L.D., Meliopoulos, V.A., Yao, J., and Schultz-Cherry, S. (2020). Obesity-Related Microenvironment Promotes Emergence of Virulent Influenza Virus Strains. MBio 11. 10.1128/mBio.03341-19.

8. Meliopoulos, V., Livingston, B., Van de Velde, L.-A., Honce, R., and Schultz-Cherry, S. (2019). Absence of β6 integrin reduces influenza disease severity in highly susceptible obese mice. J Virol 93. 10.1128/JVI.01646-18.

9. Meliopoulos, V., Honce, R., Livingston, B., Hargest, V., Freiden, P., Lazure, L., Brigleb, P.H., Karlsson, E., Sheppard, H., Allen, E.K., et al. (2024). Diet-induced obesity affects influenza disease severity and transmission dynamics in ferrets. Science Advances 10, eadk9137. 10.1126/sciadv.adk9137.

10. Schultz-Cherry, S. (2015). Role of NK cells in influenza infection. Curr Top Microbiol Immunol 386, 109–120. 10.1007/82_2014_403.

11. Moore, T., Bennett, M., and Kumar, V. (1995). Transplantable NK cell progenitors in murine bone marrow. J Immunol 154, 1653–1663.

12. Akashi, K., Traver, D., Kondo, M., and Weissman, I.L. (1999). Lymphoid development from hematopoietic stem cells. Int J Hematol 69, 217–226.

13. Fathman, J.W., Bhattacharya, D., Inlay, M.A., Seita, J., Karsunky, H., and Weissman, I.L. (2011). Identification of the earliest natural killer cell-committed progenitor in murine bone marrow. Blood 118, 5439–5447. 10.1182/blood-2011-04-348912.

14. Carlin, L.E., Hemann, E.A., Zacharias, Z.R., Heusel, J.W., and Legge, K.L. (2018). Natural Killer Cell Recruitment to the Lung During Influenza A Virus Infection Is Dependent on CXCR3, CCR5, and Virus Exposure Dose. Front Immunol 9, 781. 10.3389/fimmu.2018.00781.

15. Frank, K., and Paust, S. (2020). Dynamic natural killer cell and T cell responses to influenza infection. Front Cell Infect Microbiol 10, 425. 10.3389/fcimb.2020.00425.

16. Ge, M.Q., Ho, A.W.S., Tang, Y., Wong, K.H.S., Chua, B.Y.L., Gasser, S., and Kemeny, D.M. (2012). NK cells regulate CD8+ T cell priming and dendritic cell migration during influenza A infection by IFN-γ and perforin-dependent mechanisms. J Immunol 189, 2099–2109. 10.4049/jimmunol.1103474.

17. Gerosa, F., Baldani-Guerra, B., Nisii, C., Marchesini, V., Carra, G., and Trinchieri, G. (2002). Reciprocal activating interaction between natural killer cells and dendritic cells. J Exp Med 195, 327–333. 10.1084/jem.20010938.

18. Sole-Violan, J., Sologuren, I., Betancor, E., Zhang, S., Pérez, C., Herrera-Ramos, E., Martínez-Saavedra, M., López-Rodríguez, M., Pestano, J., Ruiz-Hernández, J., et al. (2013). Lethal influenza virus A H1N1 infection in two relatives with autosomal dominant GATA-2 deficiency. Crit Care 17, P15. 10.1186/cc11953.

19. Bongen, E., Vallania, F., Utz, P.J., and Khatri, P. (2018). KLRD1-expressing natural killer cells predict influenza susceptibility. Genome Med 10, 45. 10.1186/s13073-018-0554-1.

20. O’Shea, D., and Hogan, A.E. (2019). Dysregulation of natural killer cells in obesity. Cancers (Basel) 11. 10.3390/cancers11040573.

21. Bähr, I., Spielmann, J., Quandt, D., and Kielstein, H. (2020). Obesity-Associated Alterations of Natural Killer Cells and Immunosurveillance of Cancer. Front Immunol 11, 245. 10.3389/fimmu.2020.00245.

22. Gardiner, C.M., and Finlay, D.K. (2017). What fuels natural killers? metabolism and NK cell responses. Front Immunol 8, 367. 10.3389/fimmu.2017.00367.

23. O’Brien, K.L., and Finlay, D.K. (2019). Immunometabolism and natural killer cell\ responses. Nat Rev Immunol 19, 282–290. 10.1038/s41577-019-0139-2.

24. Keating, S.E., Zaiatz-Bittencourt, V., Loftus, R.M., Keane, C., Brennan, K., Finlay, D.K., and Gardiner, C.M. (2016). Metabolic Reprogramming Supports IFN-γ Production by CD56bright NK Cells. J Immunol 196, 2552–2560. 10.4049/jimmunol.1501783.

25. Schimmer, S., Mittermüller, D., Werner, T., Görs, P.E., Meckelmann, S.W., Finlay, D.K., Dittmer, U., and Littwitz-Salomon, E. (2023). Fatty acids are crucial to fuel NK cells upon acute retrovirus infection. Front Immunol 14, 1296355. 10.3389/fimmu.2023.1296355.

26. Kumar, P., Thakar, M.S., Ouyang, W., and Malarkannan, S. (2013). IL-22 from conventional NK cells is epithelial regenerative and inflammation protective during influenza infection. Mucosal Immunol 6, 69–82. 10.1038/mi.2012.49.

27. Nogusa, S., Ritz, B.W., Kassim, S.H., Jennings, S.R., and Gardner, E.M. (2008). Characterization of age-related changes in natural killer cells during primary influenza infection in mice. Mech Ageing Dev 129, 223–230. 10.1016/j.mad.2008.01.003.

28. Honce, R., Vazquez-Pagan, A., Livingston, B., Mandarano, A.H., Wilander, B.A., Cherry, S., Hargest, V., Sharp, B., Brigleb, P.H., Kirkpatrick Roubidoux, E., et al. (2024). Diet switch pre-vaccination improves immune response and metabolic status in formerly obese mice. Nat Microbiol 9, 1593–1606. 10.1038/s41564-024-01677-y.

29. Zhang, J., Marotel, M., Fauteux-Daniel, S., Mathieu, A.-L., Viel, S., Marçais, A., and Walzer, T. (2018). T-bet and Eomes govern differentiation and function of mouse and human NK cells and ILC1. Eur J Immunol 48, 738–750. 10.1002/eji.201747299.

30. Wang, D., and Malarkannan, S. (2020). Transcriptional regulation of natural killer cell development and functions. Cancers (Basel) 12. 10.3390/cancers12061591.

31. Littwitz-Salomon, E., Moreira, D., Frost, J.N., Choi, C., Liou, K.T., Ahern, D.K., O’Shaughnessy, S., Wagner, B., Biron, C.A., Drakesmith, H., et al. (2021). Metabolic requirements of NK cells during the acute response against retroviral infection. Nat Commun 12, 5376. 10.1038/s41467-021-25715-z.

32. Shaikh, S.R., MacIver, N.J., and Beck, M.A. (2022). Obesity dysregulates the immune response to influenza infection and vaccination through metabolic and inflammatory mechanisms. Annu Rev Nutr 42, 67–89. 10.1146/annurev-nutr-062320-115937.

33. Chandrasekaran, R., Morris, C.R., Butzirus, I.M., Mark, Z.F., Kumar, A., Souza De Lima, D., Daphtary, N., Aliyeva, M., Poynter, M.E., Anathy, V., et al. (2023). Obesity exacerbates influenza-induced respiratory disease via the arachidonic acid-p38 MAPK pathway. Front Pharmacol 14, 1248873. 10.3389/fphar.2023.1248873.

34. Alarcon, P.C., Ulanowicz, C.J., Damen, M.S.M.A., Eom, J., Sawada, K., Chung, H., Alahakoon, T., Oates, J.R., Wayland, J.L., Stankiewicz, T.E., et al. (2025). Obesity Uncovers the Presence of Inflammatory Lung Macrophage Subsets With an Adipose Tissue Transcriptomic Signature in Influenza Virus Infection. J Infect Dis 231, e317–e327. 10.1093/infdis/jiae535.

35. O’Brien, K.L., Assmann, N., O’Connor, E., Keane, C., Walls, J., Choi, C., Oefner, P.J., Gardiner, C.M., Dettmer, K., and Finlay, D.K. (2021). De novo polyamine synthesis supports metabolic and functional responses in activated murine NK cells. Eur J Immunol 51, 91–102. 10.1002/eji.202048784.

36. Terrén, I., Orrantia, A., Vitallé, J., Zenarruzabeitia, O., and Borrego, F. (2019). NK cell metabolism and tumor microenvironment. Front Immunol 10, 2278. 10.3389/fimmu.2019.02278.

37. Viel, S., Besson, L., Marotel, M., Walzer, T., and Marçais, A. (2017). Regulation of mTOR, Metabolic Fitness, and Effector Functions by Cytokines in Natural Killer Cells. Cancers (Basel) 9. 10.3390/cancers9100132.

38. Assmann, N., O’Brien, K.L., Donnelly, R.P., Dyck, L., Zaiatz-Bittencourt, V., Loftus, R.M., Heinrich, P., Oefner, P.J., Lynch, L., Gardiner, C.M., et al. (2017). Srebp-controlled glucose metabolism is essential for NK cell functional responses. Nat Immunol 18, 1197–1206. 10.1038/ni.3838.

39. Loftus, R.M., Assmann, N., Kedia-Mehta, N., O’Brien, K.L., Garcia, A., Gillespie, C., Hukelmann, J.L., Oefner, P.J., Lamond, A.I., Gardiner, C.M., et al. (2018). Amino acid-dependent cMyc expression is essential for NK cell metabolic and functional responses in mice. Nat Commun 9, 2341. 10.1038/s41467-018-04719-2.

40. Poznanski, S.M., Barra, N.G., Ashkar, A.A., and Schertzer, J.D. (2018). Immunometabolism of T cells and NK cells: metabolic control of effector and regulatory function. Inflamm. Res. 67, 813–828. 10.1007/s00011-018-1174-3.

41. Osthus, R.C., Shim, H., Kim, S., Li, Q., Reddy, R., Mukherjee, M., Xu, Y., Wonsey, D., Lee, L.A., and Dang, C.V. (2000). Deregulation of Glucose Transporter 1 and Glycolytic Gene Expression by c-Myc*. Journal of Biological Chemistry 275, 21797–21800. 10.1074/jbc.C000023200.

42. Marchingo, J.M., Sinclair, L.V., Howden, A.J., and Cantrell, D.A. Quantitative analysis of how Myc controls T cell proteomes and metabolic pathways during T cell activation. eLife 9, e53725. 10.7554/eLife.53725.

43. Dong, Y., Tu, R., Liu, H., and Qing, G. (2020). Regulation of cancer cell metabolism: oncogenic MYC in the driver’s seat. Signal Transduct Target Ther 5, 124. 10.1038/s41392-020-00235-2.

44. Marçais, A., and Walzer, T. (2014). mTOR: A gate to NK cell maturation and activation. Cell Cycle 13, 3315–3316. 10.4161/15384101.2014.972919.

45. Donnelly, R.P., Loftus, R.M., Keating, S.E., Liou, K.T., Biron, C.A., Gardiner, C.M., and Finlay, D.K. (2014). mTORC1-dependent metabolic reprogramming is a prerequisite for NK cell effector function. J Immunol 193, 4477–4484. 10.4049/jimmunol.1401558.

46. Viel, S., Marçais, A., Guimaraes, F.S.-F., Loftus, R., Rabilloud, J., Grau, M., Degouve, S., Djebali, S., Sanlaville, A., Charrier, E., et al. (2016). TGF-β inhibits the activation and functions of NK cells by repressing the mTOR pathway. Sci Signal 9, ra19. 10.1126/scisignal.aad1884.

47. Sohn, H., and Cooper, M.A. (2023). Metabolic regulation of NK cell function: implications for immunotherapy. Immunometabolism (Cobham) 5, e00020. 10.1097/IN9.0000000000000020.

48. Wang, Z., Guan, D., Wang, S., Chai, L.Y.A., Xu, S., and Lam, K.-P. (2020). Glycolysis and Oxidative Phosphorylation Play Critical Roles in Natural Killer Cell Receptor-Mediated Natural Killer Cell Functions. Front Immunol 11, 202. 10.3389/fimmu.2020.00202.

49. Burgy, O., Loriod, S., Beltramo, G., and Bonniaud, P. (2022). Extracellular Lipids in the Lung and Their Role in Pulmonary Fibrosis. Cells 11, 1209. 10.3390/cells11071209.

50. Cañadas, O., Olmeda, B., Alonso, A., and Pérez-Gil, J. (2020). Lipid–Protein and Protein–Protein Interactions in the Pulmonary Surfactant System and Their Role in Lung Homeostasis. Int J Mol Sci 21, 3708. 10.3390/ijms21103708.

51. Agassandian, M., and Mallampalli, R.K. (2013). Surfactant phospholipid metabolism. Biochim Biophys Acta 1831, 612–625. 10.1016/j.bbalip.2012.09.010.

52. Wang, B., and Tontonoz, P. (2019). Phospholipid Remodeling in Physiology and Disease. Annu Rev Physiol 81, 165–188. 10.1146/annurev-physiol-020518-114444.

53. Rong, X., Albert, C.J., Hong, C., Duerr, M.A., Chamberlain, B.T., Tarling, E.J., Ito, A., Gao, J., Wang, B., Edwards, P.A., et al. (2013). LXRs regulate ER stress and inflammation through dynamic modulation of membrane phospholipid composition. Cell Metab 18, 685–697. 10.1016/j.cmet.2013.10.002.

54. Chai, Z., Zhou, Y., Hossain, S., Huynh, K., Hajimirzaei, S., Zhang, Y., Qi, J., Zhang, G., Li, Z., and Wei, Y. Cardiolipin’s multifaceted role in immune response: a focus on interacting proteins. Front Immunol 16, 1680326. 10.3389/fimmu.2025.1680326.

55. Sheppard, S., Srpan, K., Lin, W., Lee, M., Delconte, R.B., Owyong, M., Carmeliet, P., Davis, D.M., Xavier, J.B., Hsu, K.C., et al. (2024). Fatty acid oxidation fuels natural killer cell responses against infection and cancer. Proc Natl Acad Sci USA 121, e2319254121. 10.1073/pnas.2319254121.

56. Niavarani, S.R., Lawson, C., Bakos, O., Boudaud, M., Batenchuk, C., Rouleau, S., and Tai, L.-H. (2019). Lipid accumulation impairs natural killer cell cytotoxicity and tumor control in the postoperative period. BMC Cancer 19, 823. 10.1186/s12885-019-6045-y.

57. Slattery, K., Yao, C.-H., Mylod, E., Scanlan, J., Scott, B., Crowley, J.P., McGowan, O., McManus, G., Brennan, M., O’Brien, K., et al. (2025). Uptake of lipids from ascites drives NK cell metabolic dysfunction in ovarian cancer. Science Immunology 10, eadr4795. 10.1126/sciimmunol.adr4795.

58. Kobayashi, T., Lam, P.Y., Jiang, H., Bednarska, K., Gloury, R., Murigneux, V., Tay, J., Jacquelot, N., Li, R., Tuong, Z.K., et al. (2020). Increased lipid metabolism impairs NK cell function and mediates adaptation to the lymphoma environment. Blood 136, 3004–3017. 10.1182/blood.2020005602.

59. Michelet, X., Dyck, L., Hogan, A., Loftus, R.M., Duquette, D., Wei, K., Beyaz, S., Tavakkoli, A., Foley, C., Donnelly, R., et al. (2018). Metabolic reprogramming of natural killer cells in obesity limits antitumor responses. Nat Immunol 19, 1330–1340. 10.1038/s41590-018-0251-7.

60. Gregorova, M., Santopaolo, M., Garner, L.C., Hayati, R.F., Diamond, D., Ramamurthy, N., Tran, V.T., Nguyen, N.M., Heesom, K.J., Nguyen, V.L., et al. (2025). Early NK-cell and T-cell dysfunction marks progression to severe dengue in patients with obesity and healthy weight. Nat Commun 16, 5569. 10.1038/s41467-025-60941-9.

61. Tobin, L.M., Mavinkurve, M., Carolan, E., Kinlen, D., O’Brien, E.C., Little, M.A., Finlay, D.K., Cody, D., Hogan, A.E., and O’Shea, D. (2017). NK cells in childhood obesity are activated, metabolically stressed, and functionally deficient. JCI Insight 2. 10.1172/jci.insight.94939.

62. Alarcon, P.C., Ulanowicz, C.J., Damen, M.S.M.A., Eom, J., Sawada, K., Chung, H., Alahakoon, T., Oates, J.R., Wayland, J.L., Stankiewicz, T.E., et al. (2025). Obesity Uncovers the Presence of Inflammatory Lung Macrophage Subsets With an Adipose Tissue Transcriptomic Signature in Influenza Virus Infection. J Infect Dis 231, e317–e327. 10.1093/infdis/jiae535.

63. Smith, A.G., Sheridan, P.A., Harp, J.B., and Beck, M.A. (2007). Diet-Induced Obese Mice Have Increased Mortality and Altered Immune Responses When Infected with Influenza Virus12. The Journal of Nutrition 137, 1236–1243. 10.1093/jn/137.5.1236.

64. Waggoner, S.N., Reighard, S.D., Gyurova, I.E., Cranert, S.A., Mahl, S.E., Karmele, E.P., McNally, J.P., Moran, M.T., Brooks, T.R., Yaqoob, F., et al. (2016). Roles of natural killer cells in antiviral immunity. Curr Opin Virol 16, 15–23. 10.1016/j.coviro.2015.10.008.

65. Shaikh, S.R., and Edidin, M. (2006). Polyunsaturated fatty acids, membrane organization, T cells, and antigen presentation. Am J Clin Nutr 84, 1277–1289. 10.1093/ajcn/84.6.1277.

66. Sugi, T., Katoh, Y., Ikeda, T., Seta, D., Iwata, T., Nishio, H., Sugawara, M., Kato, D., Katoh, K., Kawana, K., et al. (2024). SCD1 inhibition enhances the effector functions of CD8+ T cells via ACAT1-dependent reduction of esterified cholesterol. Cancer Science 115, 48–58. 10.1111/cas.15999.

67. von Hegedus, J.H., de Jong, A.J., Hoekstra, A.T., Spronsen, E., Zhu, W., Cabukusta, B., Kwekkeboom, J.C., Heijink, M., Bos, E., Berkers, C.R., et al. (2024). Oleic acid enhances proliferation and calcium mobilization of CD3/CD28 activated CD4+ T cells through incorporation into membrane lipids. Eur J Immunol 54, e2350685. 10.1002/eji.202350685.

68. Yang, M., Luo, S., Yang, J., Chen, W., He, L., Liu, D., Zhao, L., and Wang, X. (2022). Lipid droplet - mitochondria coupling: A novel lipid metabolism regulatory hub in diabetic nephropathy. Front. Endocrinol. 13. 10.3389/fendo.2022.1017387.

69. Fan, H., and Tan, Y. (2024). Lipid Droplet–Mitochondria Contacts in Health and Disease. Int J Mol Sci 25, 6878. 10.3390/ijms25136878.

70. Gardiner, C.M., and Finlay, D.K. (2017). What Fuels Natural Killers? Metabolism and NK Cell Responses. Front. Immunol. 8. 10.3389/fimmu.2017.00367.

71. Reed, L.J., and Muench, H. (1938). A SIMPLE METHOD OF ESTIMATING FIFTY PER CENT ENDPOINTS12. Am J Epidemiol 27, 493–497. 10.1093/oxfordjournals.aje.a118408.

72. Bligh, E.G., and Dyer, W.J. (1959). A rapid method of total lipid extraction and purification. Can J Biochem Physiol 37, 911–917. 10.1139/o59-099.

73. Palmer, A., Phapale, P., Chernyavsky, I., Lavigne, R., Fay, D., Tarasov, A., Kovalev, V., Fuchser, J., Nikolenko, S., Pineau, C., et al. (2017). FDR-controlled metabolite annotation for high-resolution imaging mass spectrometry. Nat Methods 14, 57–60. 10.1038/nmeth.4072.

